# Comparison of Synergy Extrapolation and Static Optimization for Estimating Multiple Unmeasured Muscle Activations during Walking

**DOI:** 10.1101/2024.03.03.583228

**Authors:** Ao Di, J. Fregly Benjamin

## Abstract

**Background:** Calibrated electromyography (EMG)-driven musculoskeletal models can provide great insight into internal quantities (e.g., muscle forces) that are difficult or impossible to measure experimentally. However, the need for EMG data from all involved muscles presents a significant barrier to the widespread application of EMG-driven modeling methods. Synergy extrapolation (SynX) is a computational method that can estimate a single missing EMG signal with reasonable accuracy during the EMG-driven model calibration process, yet its performance in estimating a larger number of missing EMG signals remains unclear.

**Methods:** This study assessed the accuracy with which SynX can use eight measured EMG signals to estimate muscle activations and forces associated with eight missing EMG signals in the same leg during walking while simultaneously performing EMG-driven model calibration. Experimental gait data collected from two individuals post-stroke, including 16 channels of EMG data per leg, were used to calibrate an EMG-driven musculoskeletal model, providing “gold standard” muscle activations and forces for evaluation purposes. SynX was then used to predict the muscle activations and forces associated with the eight missing EMG signals while simultaneously calibrating EMG-driven model parameter values. Due to its widespread use, static optimization (SO) was also utilized to estimate the same muscle activations and forces. Estimation accuracy for SynX and SO was evaluated using root mean square errors (RMSE) to quantify amplitude errors and correlation coefficient *r* values to quantify shape similarity, each calculated with respect to “gold standard” muscle activations and forces.

**Results:** On average, SynX produced significantly more accurate amplitude and shape estimates for unmeasured muscle activations (RMSE 0.08 vs. 0.15, *r* value 0.55 vs. 0.12) and forces (RMSE 101.3 N vs. 174.4 N, *r* value 0.53 vs. 0.07) compared to SO. SynX yielded calibrated Hill-type muscle-tendon model parameter values for all muscles and activation dynamics model parameter values for measured muscles that were similar to “gold standard” calibrated model parameter values.

**Conclusions:** These findings suggest that SynX could make it possible to calibrate EMG-driven musculoskeletal models for all important lower-extremity muscles with as few as eight carefully chosen EMG signals and eventually contribute to the design of personalized rehabilitation and surgical interventions for mobility impairments.

## Background

Muscle forces are essential for maintaining body posture and engaging in functional activities. Knowledge of the forces exerted by individual muscles is crucial for understanding the internal biomechanical mechanisms and motor control involved in human movement [1–3]. More importantly, knowledge of muscle forces could be useful for identifying musculoskeletal pathologies [4,5] and neurological disorders [6,7] as well as for designing effective rehabilitation or surgical interventions [8–10]. However, unlike joint moments, which can be measured *in vivo* directly using dynamometers or indirectly using inverse dynamics, muscle forces cannot currently be measured easily *in vivo*, though ongoing research is seeking to develop new experimental methods that can measure muscle or tendon forces *in vivo* during human movement [11,12]. Unfortunately, these research efforts have been hindered by technical challenges, high cost, and ethical considerations[11,12], motivating the search for computational methods that can enhance our knowledge of muscle forces.

Musculoskeletal modeling enables computational estimation of unmeasurable or difficult to measure internal biomechanical quantities, such as muscle forces and joint contact forces, that influence human movement generation. The estimation process uses musculoskeletal computer models that represent the bones, muscles, joints, neural control, and external forces specific to the subject and task being modeled [13–15]. These computer models typically employ a geometric model of the musculoskeletal system actuated by Hill-type muscle-tendon models [16]. The control inputs to these muscle-tendon models are either muscle excitations, which are equivalent to processed experimental electromyographic (EMG) data, or muscle activations, which are muscle excitations that have been time delayed and passed through an activation dynamics model[17]. In addition to estimating unmeasurable time-varying internal quantities (e.g., muscle activations and forces), musculoskeletal modeling can be used to estimate unmeasurable time-invariant model parameter values (e.g., optimal muscle fiber length, tendon slack length) that have a significant influence on muscle force generation [18].

The two computational methods most commonly employed for estimating muscle activations and forces using a musculoskeletal model are EMG-driven modeling [6,19–25] and static optimization (SO) [26–32]. Both methods utilize nonlinear optimization to resolve the “muscle redundancy problem” [33] (i.e., many more muscles than degrees of freedom (DOFs) in the skeleton, resulting in control indeterminacy), both require experimental joint kinematics and moments as inputs, and both find muscle activations and forces such that predicted net joint moments from a musculoskeletal model match experimental net joint moments calculated via inverse dynamics as closely as possible. However, the optimization problem formulations for these two methods are quite different (Table 1). For EMG-driven modeling, the design variables are time-invariant model parameter values (i.e., EMG scale factors, electromechanical delays, activation dynamics parameter values, Hill-type muscle-tendon model parameter values), the cost function minimizes the sum of squares of errors between model and experimental joint moments, the constraints bound muscle activations to be less than or equal to one, and the optimization problem is solved over all time frames together. For SO, the design variables are time-varying muscle activations, the cost function typically minimizes the sum of squares of muscle activations [26,34], the constraints enforce no errors between model and experimental joint moments in addition to bounds on muscle activations, and the optimization problem is solved for each time frame separately.

**Table 1:**
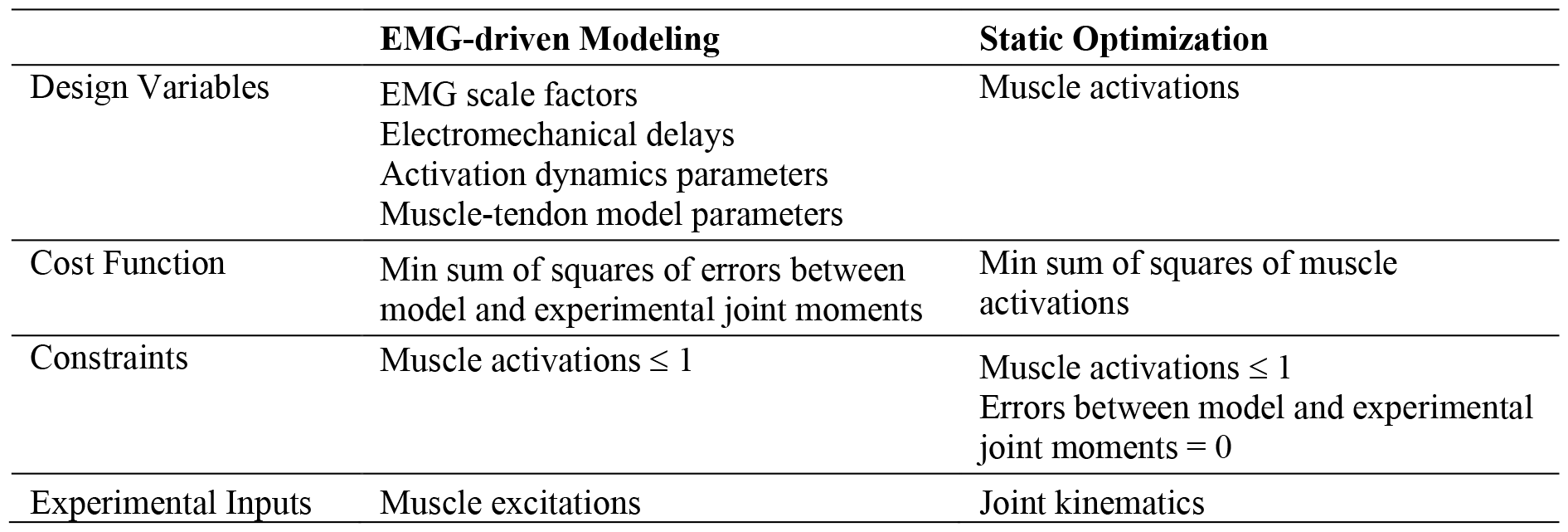
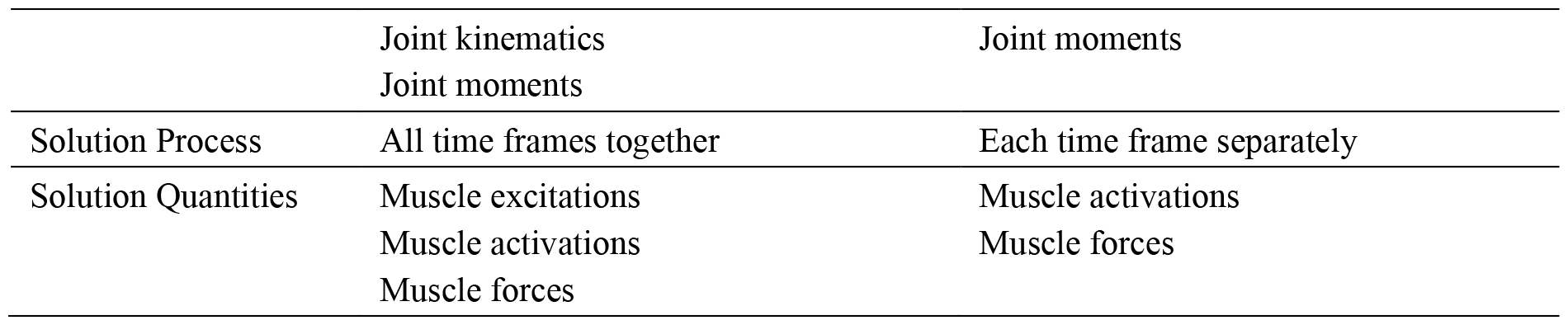
Comparison of optimization problem formulations and solutions for EMG-driven modeling and static optimization in their most fundamental forms.

**Table 2.**
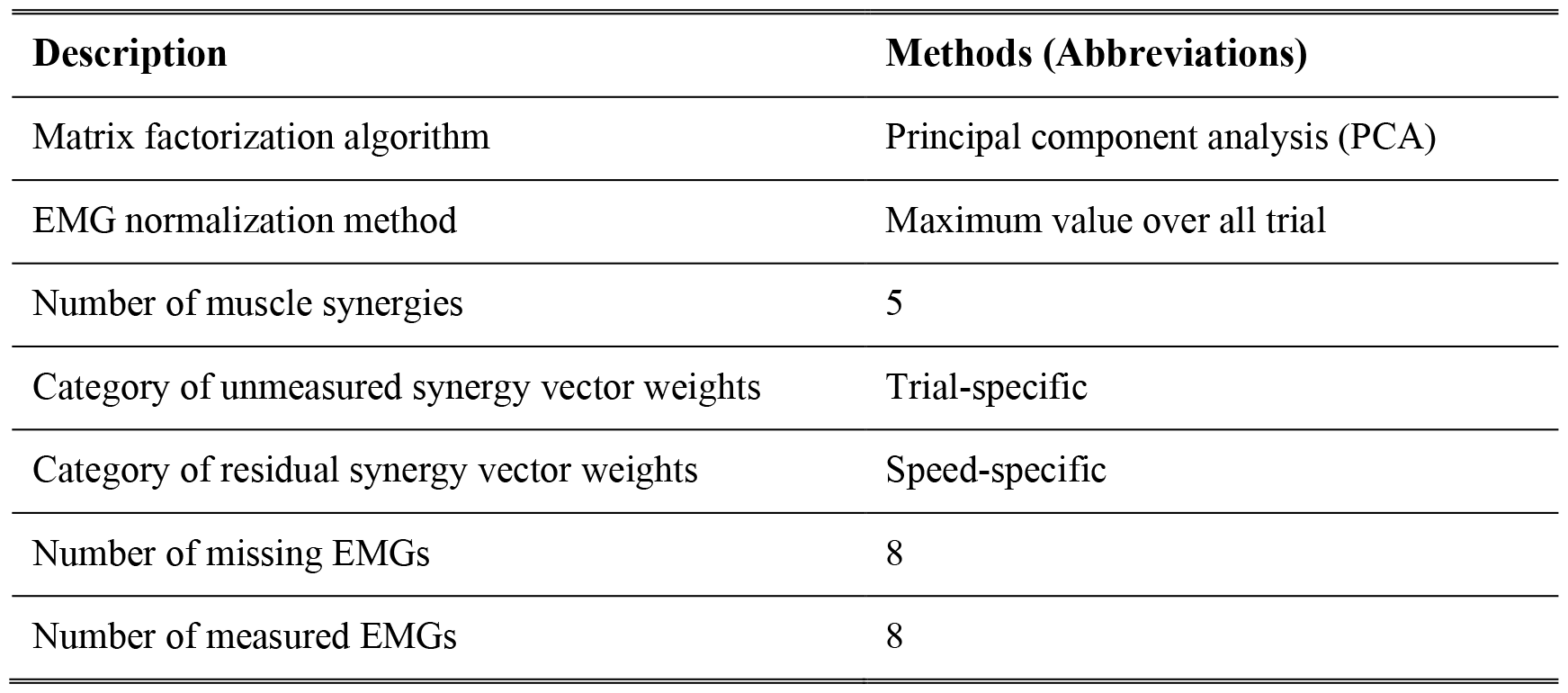
Methodological choices for synergy extrapolation.

| Description | Methods (Abbreviations) |
| --- | --- |
| Matrix factorization algorithm | Principal component analysis (PCA) |
| EMG normalization method | Maximum value over all trial |
| Number of muscle synergies | 5 |
| Category of unmeasured synergy vector weights | Trial-specific |
| Category of residual synergy vector weights | Speed-specific |
| Number of missing EMGs | 8 |
| Number of measured EMGs | 8 |

These optimization problem formulation differences, which stem from using experimental muscle excitations as inputs for EMG-driven modeling but not SO, have important implications for the capabilities and limitations of both methods. Since EMG-driven modeling uses experimental muscle excitations to constrain the time-varying shapes (and often amplitudes) of the predicted muscle excitations, model joint moments never match experimental joint moments perfectly. Consequently, minimization of this joint moment mismatch allows for calibration of musculoskeletal model parameter values when the optimization is performed over all time frames together. In contrast, since SO finds muscle activations that make model joint moments match experimental joint moments perfectly, there are no joint moment errors that can be used for calibrating musculoskeletal model parameter values. Furthermore, optimization of each time frame separately can sometimes produce muscle activation discontinuities between time frames [14,15], while minimization of muscle activations with no constraints on the time-varying shapes of the predicted muscle activations produces the smallest possible muscle activations, resulting in minimum co-contraction solutions [27,31] that may not be physiologically realistic for some subjects or movement conditions. Nonetheless, because of its simplicity and the ease with which it can be implemented and performed, SO remains the most commonly used computational method for estimating muscle activations and forces.

Although EMG-driven modeling possesses the advantages noted above and produces physiologically reasonable estimates of muscle activations and forces [24], missing EMG data from muscles that contribute significantly to a measured movement has limited the adoption of EMG-driven modeling for routine clinical gait analysis and biomechanical research. This issue is the result of two practical challenges. First, surface electrodes are incapable of measuring EMG signals from deep muscles. Despite their non-invasive nature and easy application, surface electrodes are unable to measure EMG signals from important deep muscles that contribute significantly to joint moments, such as the iliacus and psoas muscles during walking. While fine wire electrodes can capture EMG signals from deep muscles, their invasive nature, the need for specialized insertion skills, the substantial preparation time required for insertion, and the potential for discomfort and pain to the subject have limited their utilization. Furthermore, in certain scenarios, deep muscles may be inaccessible even with fine wire electrodes. For instance, the use of a fine wire electrode is contraindicated for safety reasons in subjects with a cancerous tumor near the muscle to be measured. Second, EMG systems possess a limited number of channels for collecting EMG data. Many EMG systems available in human movement labs provide support for 16 channels of data, which means only eight channels of EMG data can be collected per leg when measuring activities such as walking or running. However, EMG-driven lower extremity models close to 16 channels per leg to inform the model without omitting any important large muscles. These challenges are significant as the absence of EMG data from important muscles can have a negative impact on the reliability of force estimates for other muscles that span the same joints [25,29]. To address the issue of missing EMG signals when performing EMG-driven modeling, researchers either exclude muscles with missing EMG data from the musculoskeletal model [28,35], include such muscles in the model but assume that they generate only passive force [25], or include such muscles and use SO to estimate the associated muscle activations [28,29].

To provide a better alternative for addressing missing EMG signals, researchers have recently developed a modified EMG-driven modeling approach called “Synergy Extrapolation” (SynX) that uses muscle synergy concepts to estimate missing muscle excitation data [36–38]. The theoretical basis for SynX is that a large number (e.g., 8 or 16) of experimentally measured muscle excitations can be represented by a smaller number (e.g., 4 or 5) of muscle synergies composed of time-varying synergy excitations and associated time-invariant synergy vectors, where the weights in each synergy vector define how the associated synergy excitation contributes to all muscle excitations. The synergy excitations provide information about the timing of muscle contractions, while the synergy vectors provide information about the coordination of muscle contractions. Given 16 experimental muscle excitations, if a lower dimensional set of 4 or 5 muscle synergies are calculated using either all 16 excitations or a subset of 8 carefully selected excitations, the resulting synergy excitations will be almost the same in both cases [36]. This observation demonstrates that the time-varying synergy excitations extracted from the first 8 muscle excitations can be used as basis functions for constructing the remaining 8 muscle excitations.

Based on this observation, the historical development of SynX followed a logical sequence of three studies. First, SynX was shown to work in theory for fitting eight missing muscle excitations using synergy excitations extracted from eight measured muscle excitations [36]. For this study, 16 muscle excitations per leg measured experimentally from three subjects during walking were split into two groups of eight “measured” and eight “missing” excitations, and synergy excitations calculated from the eight measured excitations were used to fit the eight missing excitations. This study only established the theoretical feasibility of SynX, since the fitting process required the use of the missing muscle excitations. Second, SynX was shown to work in practice for predicting a single missing muscle excitation if a musculoskeletal model with pre-calibrated parameter values was used in the process [37]. The same sets of 16 experimental muscle excitations were again split into two groups, where 15 muscle excitations were treated as “measured” and one muscle excitation at a time collected from a fine wire electrode was treated as “missing.” A key limitation of this study was the need for a pre-existing calibrated musculoskeletal model before the missing muscle excitation could be predicted reliably, which necessitates a priori knowledge of the missing muscle excitation for initial model calibration. Third, SynX was shown to work in practice for predicting a single missing muscle excitation while simultaneously calibrating musculoskeletal model parameter values [38]. The same sets of 16 experimental muscle excitations were again split into groups of 15 “measured” muscle excitations and one “missing” fine wire muscle excitation. A multi-objective optimization problem was designed to predict the missing muscle excitation while simultaneously calibrating time-invariant musculoskeletal model parameter values and time-varying residual muscle activations needed to account for small errors in the measured muscle excitations. This study resolved the main limitation of the previous study by allowing EMG-driven model calibration and prediction of a single missing muscle excitation to be performed simultaneously. SynX has been used more recently to predict the activation of a single unmeasured upper-extremity muscle (e.g. biceps long head), achieving a Pearson’s correlation coefficient of up to 0.99 with the same muscle activation calculated from experimental EMG data withheld for evaluation purposes [39]. The next logical study in this progression is to evaluate how well SynX works in practice for predicting multiple missing muscle excitations while simultaneously calibrating musculoskeletal model parameter values. If SynX can predict missing muscle excitations reliably using a low number of EMG signals collected using only surface electrodes, the applicability of EMG-driven modeling to research and clinical questions will be greatly expanded.

This study evaluated how well SynX can estimate muscle activations associated with eight channels of missing EMG data using synergy excitations extracted from muscle excitations associated with eight channels of measured EMG data while simultaneously calibrating musculoskeletal model parameter values. Experimental walking data collected from two subjects post-stroke were used for the evaluation. Time-varying quantities (muscle activations and forces along with net joint moments) and time-invariant model parameter values (activation dynamics and Hill-type muscle-tendon model parameter values) predicted by SynX were compared to “gold standard” results produced by EMG-driven model calibration using a complete set of EMG data where no EMG signals were regarded as missing. Time-varying quantities (muscle activations and forces) predicted by SO were also compared to the “gold standard” results to determine which method provides the most reliable predictions. In addition, the reliability with which SynX and SO can predict muscle activations and forces when using pre-calibrated musculoskeletal models was evaluated to assess how model calibration affects muscle activation and force estimates from both methods.

## Methods

### Experimental Data Collection

Two previously published experimental walking datasets collected from a high-functioning hemiparetic subject (S1, male, 1.70 m tall, mass 80.5 kg, age 79 years, right side hemiparesis, lower extremity Fugl-Meyer Motor Assessment score of 32 out of 34) and a low-functioning hemiparetic subject post-stroke (S2, male, 1.83 m tall, mass 88.5 kg, age 62 years, right side hemiparesis, lower extremity Fugl-Meyer Motor Assessment score of 25 out of 34) were used for this study [23,40]. After giving written informed consent, both subjects walked on a split-belt instrumented treadmill (Bertec Corp., Columbus, OH, United States) at their self-selected speed (0.5 m/s for S1 and 0.35 m/s for S2) and fastest-comfortable speed (0.8 m/s for S1 and 0.65 m/s for S2). All experimental procedures were approved by the University of Florida Health Science Center Institutional Review Board (IRB-01).

Sixteen channels of EMG data were collected from each leg of both subjects using both surface and fine wire electrodes (Motion Lab Systems, Baton Rouge, LA, United States). These extensive EMG data enabled every muscle in each leg of each subject’s musculoskeletal model (see below) to have an associated experimental EMG signal, providing an opportunity to verify the reliability of muscle activations and forces estimated by SynX and SO. Surface EMG data were collected from the following superficial muscle groups (figure 1): 1) *GlutMax*, including gluteus maximus superior (*glmax1*), gluteus maximus middle (*glmax2*) and gluteus maximus inferior (*glmax3*); 2) *GlutMedMin*, including gluteus medius anterior (*glmed1*), gluteus medius middle (*glmed2*), gluteus medius posterior (*glmed3*), gluteus minimus anterior (*glmin1*), gluteus minimus middle (*glmin2*), and gluteus minimus posterior (*glmin3*); 3) *SemiMembTen*, including semimembranosus (*semimem*) and semitendinosus(*semiten*); 4) *RecFem*, including rectus femoris (*recfem*); 5) *Bicfem*, including biceps femoris long head (*bflh*) and biceps femoris short head (*bfsh*); 6) *VasMedInt*, including vastus medialis (*vasmed*) and vastus intermedius (*vasint*); 7) *VasLat*, including vastus lateralis (*vaslat*); 8) TibAnt, including tibialis anterior (*tibant*); 9) *Peroneus*, including peroneus brevis (perbrev) and peroneus long (perlong); 10) *Sol*, including soleus (soleus). Additionally, fine-wire EMG data were collected from the following deep muscle groups (Fig.2): 1) *iliopsoas*, including iliacus (*iliacus*) and psoas (*psoas*); 2) *Adductors*, including adductor brevis (*addbrev*), adductor longus (*addlong*), adductor magnus distal (*addmagDist*), adductor magnus ischial (*addmagIsch*), adductor magnus middle (*addmagMid*), and adductor magnus proximal (*addmagProx*); 3)*Tibpost*, including tibialis posterior (*tibpost*). Small differences existed in the EMG data collect from the two subjects. For the high-functioning subject (S1), a surface EMG signal (referred as *GasMed)* was also collected and expanded to medial gastrocnemius (*gasmed*) and lateral gastrocnemius (*gaslat*), and two fine-wire EMG signals (referred as *ExtDigLong* and *FlexDigLong*) were recorded from extensor digitorum longus (*edl*) and flexor digitorum longus (*fdl*) respectively. For the low-functioning subject (S2), two surface EMG signals (referred as *GasMed* and *GasLat*) were recorded from medial gastrocnemius (*gasmed*) and lateral gastrocnemius (*gaslat*) respectively, and a fine-wire EMG signal (referred as *TensFascLat*) was recorded from tensor fasciae latae (*tfl*). Raw EMG data were high-pass filtered at 40 Hz, demeaned, full-wave rectified, and low-pass filtered at 3.5/*tf* Hz, where where *tf* is the period of the gait cycle [23].Processed EMG data were then normalized to the maximum values across all experimental gait cycles. The resulting processed EMG data will henceforth be referred to as “experimental muscle excitations” [23,41].

**Figure 1.**
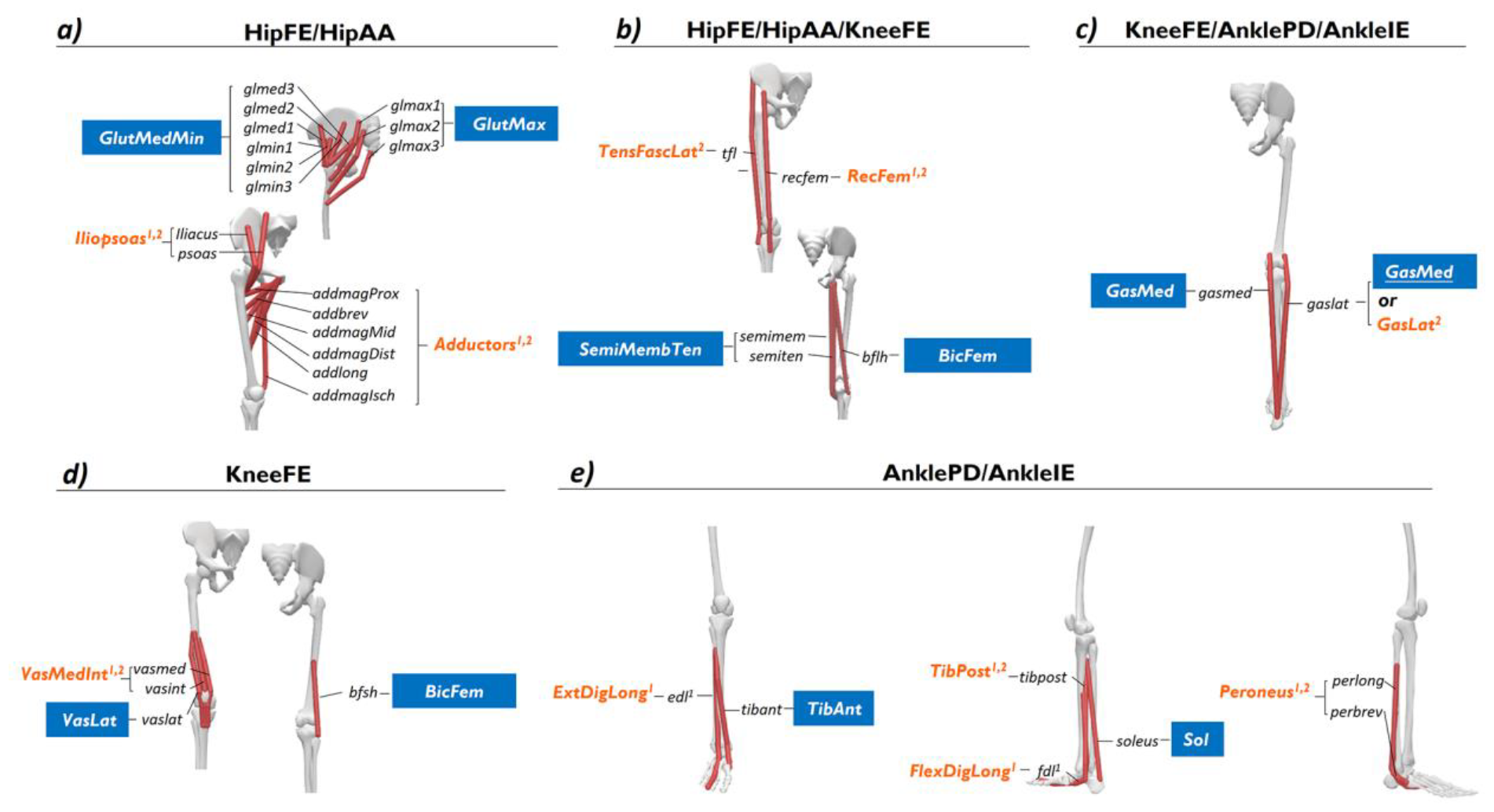
The assumption about “measured” and “unmeasured” EMG channels when performing SynX and SO as well as the associated muscles in the OpenSim model for each subject. The EMG channels assumed “measured” are denoted by blue boxes, while those assumed “unmeasured” are indicated by orange italic texts. The superscripts 1 and 2 represent the assumption of “unmeasured” EMG channels for subject S1 and S2, respectively. The muscles were categorized based on their actuating degrees of freedom (DOFs).

A three-dimensional motion capture system (Vicon Corp., Oxford, United Kingdom) operating at 100 Hz was used to measure reflective surface marker trajectories, while two treadmill force plates (Bertec Corp., Columbus, OH, United States) recording at 1000 Hz were used to measure ground reaction forces and moments. Raw motion capture and ground reaction data were low-pass filtered with a variable cut-off frequency of 7/*tf* Hz [42], where *tf* is the period of the gait cycle. Data from ten gait cycles (five cycles per speed) per leg were randomly chosen to simultaneously calibrate the EMG-driven models and evaluate the accuracy of estimated muscle activations and forces. Following pre-processing, data from each gait cycle were resampled to 101 time points from heel-strike (0%) to subsequent heel-strike (100%) of the same foot. An extra 20 time frames, accounting for a maximum electromechanical delay of approximately 100 ms, were retained prior to the start of each gait cycle, yielding 121 time points for each of the 10 gait cycles.

### Musculoskeletal Model Creation

A generic full-body OpenSim musculoskeletal model [43] was used as the starting point to create a personalized model of each subject. This generic model possessed 37 degrees of freedom (DOFs), 80 muscle-tendon actuators to control lower limb motion, and 17 ideal torque actuators to control the upper body motion. For each subject, a sequence of four analyses were performed using OpenSim 4.0 [44,45] to prepare the model for EMG-driven modeling with SynX. First, OpenSim model scaling was performed so that the generic model’s anthropometry would more closely match that of each subject. Second, repeated OpenSim inverse kinematics (IK) analyses within a nonlinear optimization were performed to calibrate the locations and orientations of lower body joint centers and axes such that errors between model and experimental surface marker positions were minimized for isolated joint motion and walking trials [46]. The lower body DOFs affected by this calibration process were hip flexion/extension (*HipFE*), hip adduction/abduction (*HipAA*), hip internal/external rotation (*HipRot*), knee flexion/extension (*KneeFE*), ankle plantarflexion/dorsiflexion (*AnklePD*), and ankle inversion/eversion (AnkleIE). These six low-extremity DOFs were targeted because their associated experimental joint moments were needed for performing SynX and SO. Third, additional OpenSim IK analyses were performed using experimental marker data from the walking trials to obtain joint angle time histories. Fourth, OpenSim inverse dynamic (ID) analyses were performed using the previously calculated joint kinematics and the experimental ground reaction data from the walking trials to calculate experimental joint moments.

### Muscle Activation and Force Estimation

SynX and SO were both utilized to estimate muscle activations and forces, and the resulting estimates from both methods were compared to a “gold standard” for evaluation purposes. As illustrated in figure 2, both approaches take joint kinematics and associated musculoskeletal geometries (i.e., muscle-tendon lengths and moment arms) as inputs to estimate muscle activations, muscle forces, and net joint moments. Subsequently, the estimated predicted net joint moments are iteratively compared to the inverse dynamic joint moments through an optimization process, leading to the estimation of the time-varying muscle activations and forces for SynX and SO, as well as time-invariant musculoskeletal model and SynX-related parameter values for SynX.

**Figure 2.**
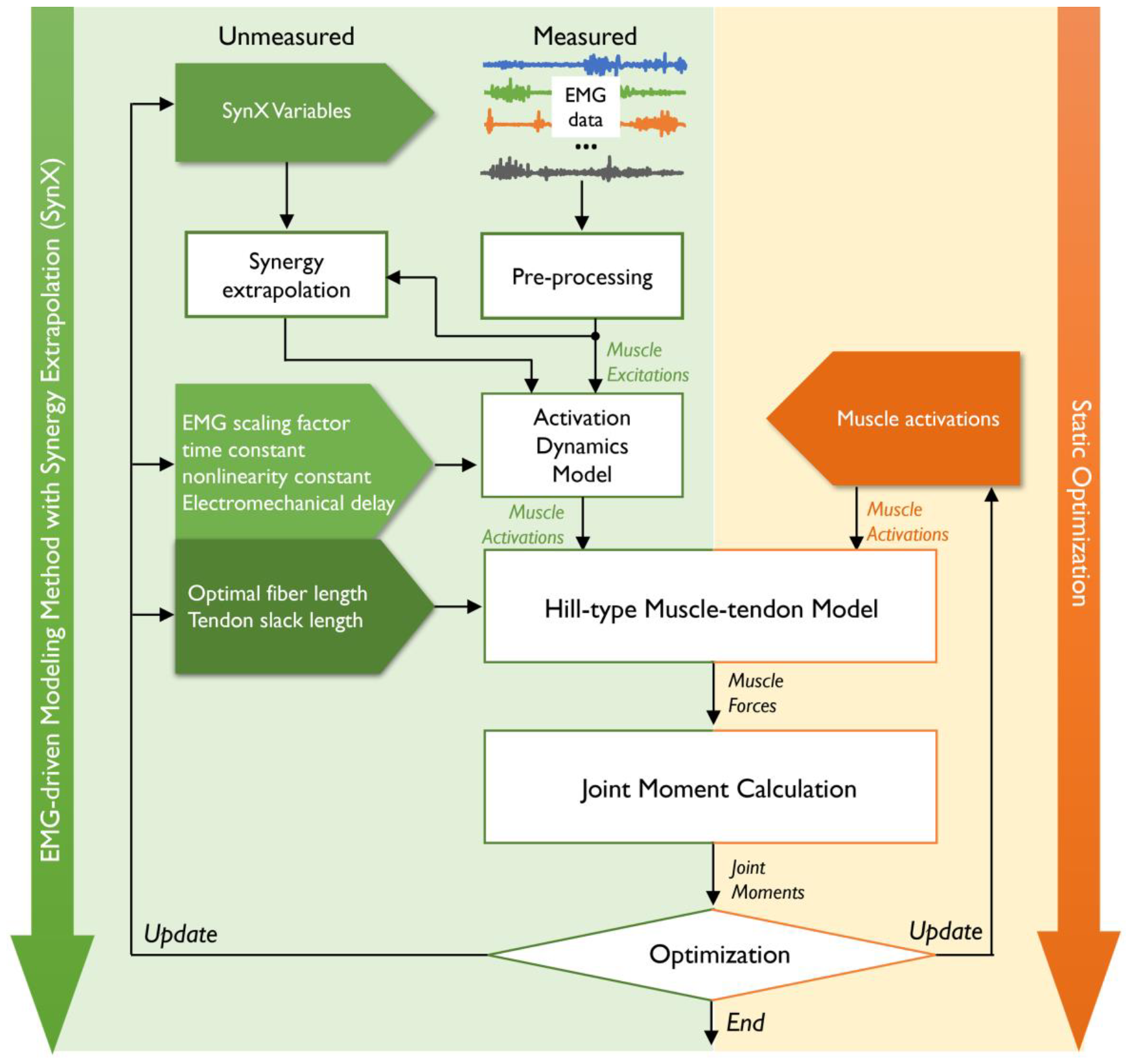
The workflow for EMG-driven modeling with SynX (left panel with a green background) and SO (right panel with an orange background) as performed in this study. Both methods employ experimental joint kinematics and moments as inputs and aim to determine muscle activations and forces in such a way that the predicted net joint moments from a musculoskeletal model closely match the experimental net joint moments calculated via inverse dynamics. However, there are notable differences in the optimization problem formulations for these two methods. In EMG-driven modeling with SynX, the design variables consist of time-invariant model parameter values and SynX variables, with the optimization problem being solved across all time frames together. Conversely, for SO, the design variables encompass time-varying muscle activations, typically utilizing model parameter values from scaled generic models or literature references, and the optimization problem is solved for each time frame separately. Subsequently, both techniques leverage the Hill-type muscle-tendon model to estimate muscle forces and their respective contributions to the joint moments.

### Synergy Extrapolation Solution Process

The SynX solution process involved four tasks as summarized below.

#### Muscle activation estimation

For the first task of the SynX solution process, muscle activations were found for muscles with and without experimental EMG data. The transformation of excitations from measured muscles into activations of all muscles itself involved four distinct steps [37,38]. First, muscle excitations 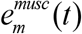 for muscles with experimental EMG data were adjusted using a muscle-specific scale factor ranging from 0.05 to 1, acknowledging that actual maximum activation levels tend to surpass those observed experimentally during walking.

Second, muscle synergy analysis (MSA) was conducted on the scaled muscle excitations using principal component analysis (PCA) to extract a small number of muscle synergies, specifically six for the present study:

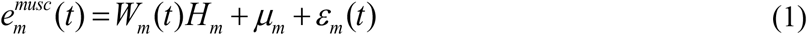

Where 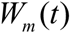 specifies the time-varying measured synergy excitations, *H*_*m*_ specifies the associated measured synergy vector weights, *μ*_*m*_ stands for the average values of each measured muscle excitation, and *ε*_*m*_ (*t*) stands for the decomposition residuals that could not be accounted for by *W*_*m*_ (*t*)*H*_*m*_ + *μ*_*m*_. Following MSA, both unmeasured muscle excitations 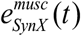 and residual muscle excitations *e*^*res*^ (*t*) added to the measured muscle excitations were constructed from the measured synergy excitations:

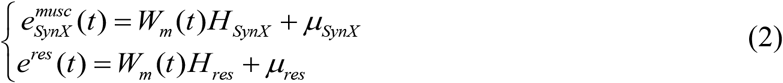

Where *H*_*SynX*_ represents the unmeasured synergy vector weights, *μ*_*SynX*_represents the average values of each unmeasured muscle excitation, *H*_*res*_ represents the residual synergy vector weights, and *μ*_*res*_ represents the average values of each residual muscle excitation. Henceforth, we denote the union of *H*_*SynX*_ , *μ*_*SynX*_ , *H*_*res*_ and *μ*_*res*_ as SynX variables, which were all time-invariant and determined through an optimization process implemented within the EMG-driven model calibration process (Figure 2). Once unmeasured and residual muscle excitations were constructed, two sets of muscle excitations were calculated when residual muscle excitations were and were not included:

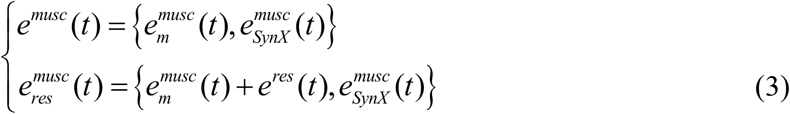

where *e*^*musc*^ (*t*) defines the muscle excitations without residual muscle excitations included, while 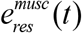 defines the muscle excitations with residual muscle excitations included. Both *e*^*musc*^ (*t*) and 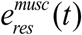 were utilized in subsequent steps to compute corresponding muscle activations denoted as *a*^*musc*^ (*t*) and 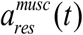, respectively.

Third, neural activations *u*^*musc*^ (*t*) were determined from constructed muscle excitations by employing a first-order ordinary differential equation for activation dynamics [47]:

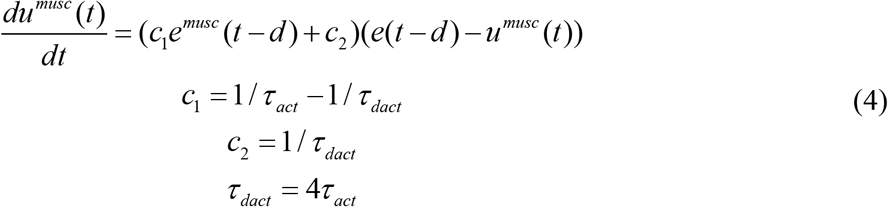

Where *τ* _*act*_ and *τ*_*dact*_ are activation and deactivation time constants. *d* specifies the electromechanical time delay.

Fourth, a nonlinear one-parameter transformation model was utilized to compute each associated muscle activation *a*^*musc*^ (*t*) [48]:

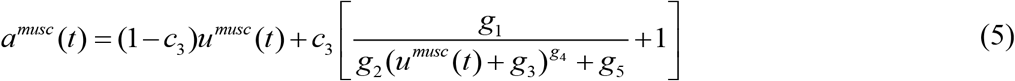

Where *c*_3_ is an activation nonlinearity constant that characterizes the curvature of the relationship of each muscle. *g*_1_ to *g*_5_ are constant coefficients obtained by fitting published experimental data from isometric contractions[48]. Our EMG-driven modeling approach solves for muscle activations with (i.e., 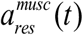) or without (i.e., *a*^*musc*^ (*t*) ) residual excitations included over all time frames simultaneously by adjusting the same set of design variables, encompassing SynX variables, EMG scale factors, electromechanical time delays, activation time constants, and activation nonlinearity constants, where further details are provided in section 2.3.1.4.

#### Muscle force estimation

For the second task of the SynX solution process, muscle forces were estimated using the activations for measured and unmeasured muscles found in the first task. Taking the estimated muscle activations as inputs, our EMG-driven modeling process employed a Hill-type muscle tendon model with rigid tendon [16,23,49] to predict the force generated by a given muscle-tendon actuator, *m* , which was formulated as (figure 2):

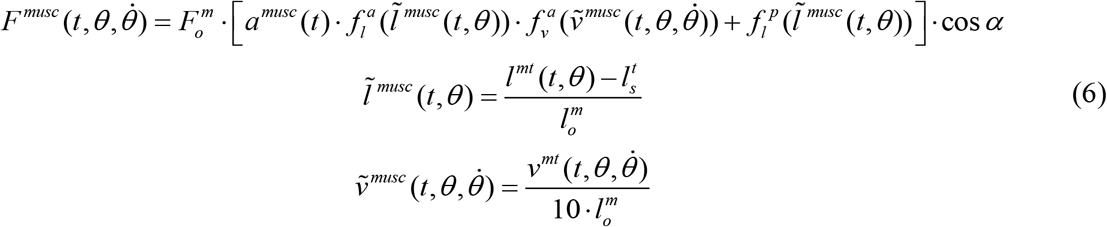

Where 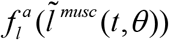 and 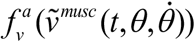 describe the normalized active muscle force-length and force-velocity relationships, respectively,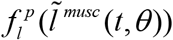 defines the normalized force-length relationship, 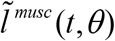 and 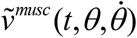 denote the time-varying normalized muscle fiber length and velocity, respectively, 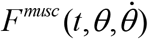 and *a*^*musc*^ (*t*) denote the muscle force and muscle activation generated by the muscle-tendon actuator at time *t* , 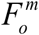 is the maximum isometric force, *α* is the pentation angle of the muscle (values of which were taken from literature [50]), 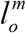 denotes optimal muscle fiber length, and 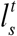 denotes tendon slack length. These values (apart from pennation angles, which were taken from the literature) were calibrated through an optimization process or taken from the scaled OpenSim models. More details regarding the determination of 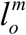 and 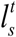 values for each muscle force estimation method can be found below.

#### Joint moment calculation

For the third task of the SynX solution process, model joint moments were calculated using the forces for measured and unmeasured muscles found in the second task. Once the muscle forces *F*^*musc*^ (*t*,*θ* ) were estimated, their contributions to net joint moment at joint *j* were calculated using the corresponding muscle moment arms:

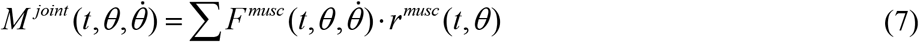

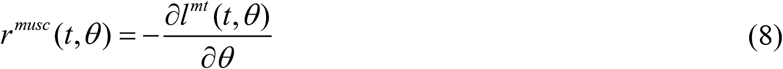

Where 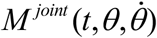 is joint moment at time *t* , which is defined as the sum of contributions from all spanning muscles *r*^*musc*^ (*t*,*θ* ) is muscle moment arm for muscle *m* at time *t* , which was defined as the negative of the partial derivative of muscle-tendon length *l*^*mt*^ (*t*,*θ*) with respect to generalized coordinate *θ* [51]. The negative sign in Eq. 8 was implemented for consistency with the OpenSim modeling environment. When utilizing SynX for estimating unmeasured muscle excitations, net joint moments were computed with 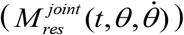and without 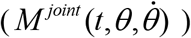 incorporating residual excitations into the measured muscle excitations, as stipulated by the cost function for EMG-driven model calibration.

#### Optimization problem formulation

For the fourth task of the SynX solution process, the first three tasks were performed iteratively within a nonlinear optimization that adjusts three categories of design variables (see figure 2): 1) SynX parameter values including synergy vector weights and average values associated with unmeasured muscle excitations as well as synergy vector weights and average values associated with residual muscle excitations; 2) activation dynamics model parameter values consisting of EMG scale factors, electromechanical delays, activation time constants, and activation nonlinearity constants; 3) muscle-tendon model parameter values consisting of optimal muscle fiber lengths and tendon slack lengths. EMG-driven model calibration typically adjusts muscle forces by altering muscle-tendon model parameter values such that the differences between model-predicted and inverse dynamic (ID) joint moments are minimized. However, to estimate unmeasured muscle excitations via SynX during EMG-driven model calibration, the primary cost function was formulated as a trade-off between minimizing joint moment tracking errors and minimizing unmeasured and residual muscle activation magnitudes [38]:

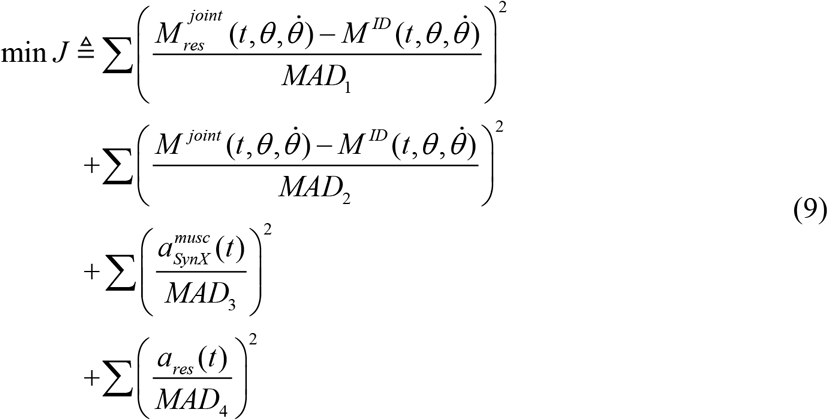

where 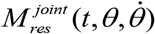 refers to model-predicted joint moments when residual muscle excitations are included in joint moment calculations, 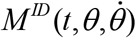 refers to inverse dynamic joint moments obtained from OpenSim ID analyses, 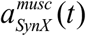 represents unmeasured muscle activations estimated by SynX, and *a*_*res*_ (*t*) signifies residual muscle activations added to the measured muscle activations, which are equivalent to 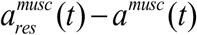. Normalization of all four cost function terms was achieved using a set of maximum allowable deviations (MAD), the values of which were determined by performing a sensitivity analysis as described in [38]. Further details regarding initial guesses, upper/lower bounds for design variables, additional inequality constraints, and penalty terms can be found in previously published studies [23,37,38]. All optimization procedures were performed using MATLAB’s ”*fmincon*” function with its sequential quadratic programming algorithm.

### Static Optimization Solution Process

The static optimization solution process involved determining muscle activations *a*^*musc*^ (*t*) at each time instant *t* by performing an inverse dynamics-based optimization. In the standard SO approach, the muscle redundancy problem is addressed by minimizing the energetic cost represented by the sum of squares of muscle activations while ensuring that inverse dynamic joint moments are matched perfectly at the solution [26]:

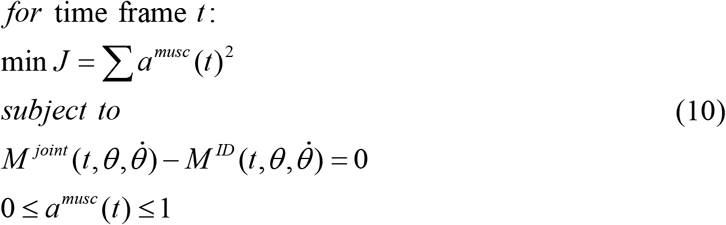

The net joint moments for SO were estimated by multiplying the muscle forces obtained through substituting muscle activations *a*^*musc*^ (*t*) into the Hill-type muscle-tendon model with the corresponding muscle moment arms, as depicted in Eqs. 6 through 8. In contrast to the EMG-driven modeling method, the muscle activations estimated for SO were used directly as design variables in the optimizations, which were solved individually for each time frame. Furthermore, model parameter values were taken from scaled generic models or literature references rather than being calibrated during the optimization process.

### Synergy Extrapolation and Static Optimization Evaluation

#### Muscle selection heuristics

Given 16 measured muscle excitations for each leg of both subjects, we had to decide which 8 muscle excitations would be treated as measured and which 8 would be held back and treated as missing for SynX and SO evaluation purposes. A prior study [36] provided guidance for which eight muscles to select as measured and which eight to select as missing so as to maximize reconstruction accuracy for the eight missing muscle excitations. In that study, an investigation of all possible combinations of eight measured and eight missing EMG signals yielded the following muscle selection heuristic: 1) Choose muscles easily accessible by surface EMG electrodes; 2) Choose most frequently occurring muscle in the top 10% of muscle combinations that yielded the highest SynX accuracy from each primary lower extremity function group; 3) Choose two hip/knee biarticular muscles at minimum; 4) Choose remaining most frequent muscles to fill eight muscle combinations. Following the observation of SynX performance, our muscle selection heuristic, given a limited number of eight EMG channels, indicated that researchers should collect surface EMG data from commonly chosen uniarticular and biarticular flexor and extensor muscles from each major muscle group, as illustrated in figure 1. The selected uniarticular muscles included a hip extensor (*GlutMax*), a knee extensor (*VasLat* considered preferable over *VasMed*), an ankle plantarflexor (*Sol*), and an ankle dorsiflexor (*TibAnt*). Uniarticular hip flexor (*Iliopsoas*) was omitted due to the difficulty in measuring these muscles reliably with surface electrodes. The chosen biarticular muscles included a posterior thigh muscle (*SemiMembTen*, or *Bicfem*), and a posterior calf muscle (*GasMed* or *GasLat*). Additionally, adding *GlutMedMin* to the list appeared to be a reasonable choice if one more muscle was needed. Even the collection of EMG data from less commonly chosen muscles spanning all three joints (*Adductors, tfl*, and *Peroneus*) may facilitate to improve the estimation accuracy to some extent due to the unique stabilizing roles they play in the frontal plane, they were excluded from the “measured” muscles that was attributed to the difficulty in reliable surface EMG measurement of these muscles.

#### Synergy extrapolation methodological choices

Implementation of SynX requires making several methodological choices that can impact the accuracy of estimated muscle activations and forces. Previous studies investigated the influence of various methodological factors on SynX performance [37,38], including EMG normalization methods, matrix decomposition algorithms, the number of muscle synergies, and assumptions regarding the variability of synergy vector weights across trials for the reconstruction of unmeasured and residual muscle excitations. We systematically assessed the results for all possible methodological combinations and found that principal component analysis (PCA) with either five or six synergies consistently predicted unmeasured muscle excitations with reasonable accuracy.

In contrast, non-negative matrix factorization (NMF) did not achieve acceptable prediction accuracy. Additionally, for the same number of synergies, employing trial-specific unmeasured synergy vector weights and speed-specific residual synergy vector weights resulted in optimal SynX performance for both subjects in terms of estimation accuracy and computational efficiency. Notably, EMG normalization had no significant impact on SynX performance. Thus, the key methodological choices for SynX in this study were informed by insights from prior research, as detailed in Table 1.

#### Optimization problems

In this study, three primary objectives were pursued. Firstly, the study aimed to evaluate the performance of SynX when treating multiple channels of EMGs (i.e., eight) as “unmeasured”. Secondly, it sought to compare the estimates of muscle activations and forces from SynX with those from SO. Thirdly, the study also aimed to analyze the accuracy of estimated unmeasured muscle activations and forces for both SynX and SO when using model parameter values associated with different levels of personalization.

To address these primary objectives, we formulated six optimization problems to estimate unmeasured muscle activations and, for SynX, calibrate model parameter values (figure 3). The first optimization problem utilized all 16 channels of EMG data to calibrate each EMG-driven musculoskeletal model, providing “gold standard” muscle activations and forces for evaluation (termed “ *Params* ”). The second optimization problem assessed the performance of SynX when multiple channels of EMG data (i.e., eight) were considered “unmeasured”. This optimization problem calibrated EMG-driven models for each leg of each subject while simultaneously estimating missing muscle excitations using SynX, where activation dynamics model, muscle-tendon model, and SynX parameter values were calibrated concurrently (termed “ *S*yn*X*_*Unmeasured*_ + *Params* ”). The third optimization problem used SO to estimate muscle activations for all muscles using muscle-tendon model parameter values taken from scaled generic OpenSim models (termed 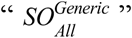 ), representing the most commonly formulated SO method. The accuracy of estimated muscle activations and forces was further quantitatively compared to those from the optimization “ *S*yn*X*_*Unmeasured*_ + *Params* ” to assess the estimation performance of both SynX and SO. The fourth optimization problem employed SynX to estimate the unmeasured muscle excitations within a well-calibrated EMG-driven model (termed 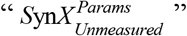 ), utilizing the model parameter values found in the “gold standard ( *Params )*” optimization. The fifth and sixth optimization problems utilized SO to estimate muscle activations for all muscles (termed 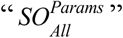) and only unmeasured muscles (termed 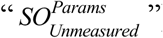) using the model parameter values from the “gold standard ( *Params )*” optimization, rather than scaled generic values. When performing the fourth and sixth optimizations of 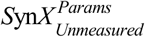 and 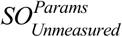 to estimate only unmeasured muscle excitations/activations, the muscle activations of the measured muscles were determined from the “gold standard ( *Params )*” optimization.

**Figure 3.**
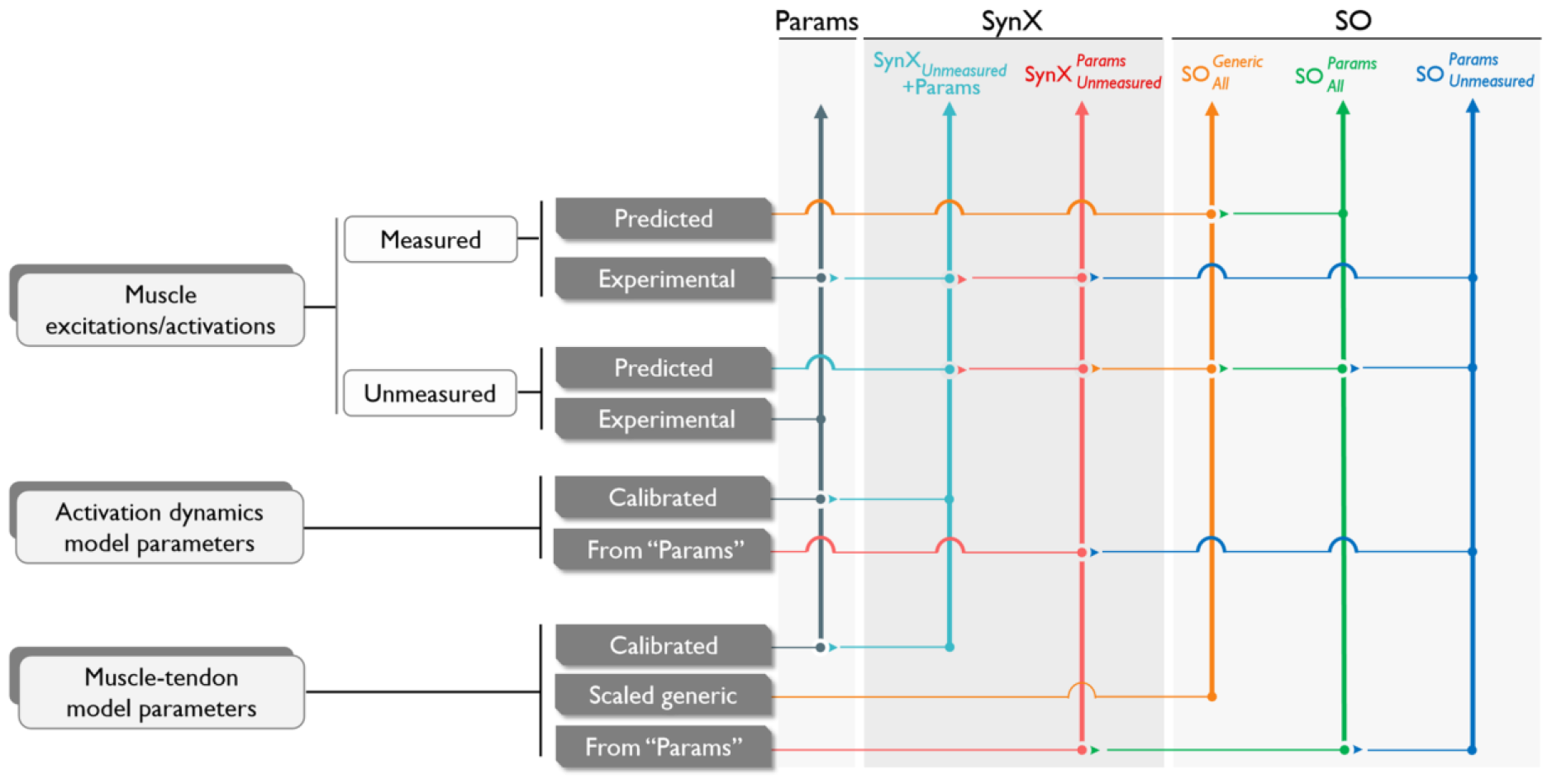
Summary of six optimizations performed in this study, which included two optimizations using SynX to predict unmeasured muscle excitations (*termed SynX Unmeasured* +*Params* and 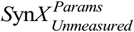), three optimizations using static optimization (SO) to predict unmeasured muscle activations (termed 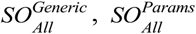 and 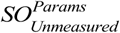) and one “gold standard” optimization using the complete set of EMG signals with no muscle excitations predicted by SynX or SO (termed *Params* ). The calibration cases were named based on the prediction method for unmeasured muscle excitations or activations as well as the categories of design variables included in the optimization problem formulation. The subscripts indicate which set of muscle excitations or activations were predicted computationally, while the superscripts indicate which set of model parameters were employed during model calibration. In each column of the optimizations, the arrows indicate whether each group of muscle excitations or activations were predicted or obtained experimentally. Moreover, the arrows indicate which values were used if the model parameters were not calibrated through optimization. The term “Scaled Generic” denotes the scaled generic model parameter values, while “From Params” refers to the model parameter values derived from the “gold standard (*Params*)” optimization.

### Evaluation Metrics and Statistical Analyses

Several common evaluation metrics were utilized to evaluate the ability of SynX and SO to estimate muscle activations and muscle forces for unmeasured muscles and joint moments across all cases. First, root mean square errors (RMSEs) were computed to quantify magnitude errors between experimental (from “ *Params* ” case) and predicted (from two SynX and three SO cases) muscle activations and forces. Similarly, Pearson correlation coefficients (*r*) were computed to quantify shape similarity between experimental and predicted unmeasured muscle activations and forces. Correlations were interpreted based on [52], categorized as weak (*r* < 0.35), moderate (0.35 < *r* ≤ 0.67), strong (0.67 < *r* ≤ 0.9), or very strong (*r* ≤ 0.9). Furthermore, mean absolute errors (MAEs) between model and experimental net joint moments were also calculated for the “*Params*” case and the two SynX cases “ *S*yn*X*_*Unmeasured*_ + *Params* ” and 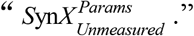 Evaluation metrics, including RMSEs, *r* values, and MAE values, were calculated by concatenating the data across all calibration trials and legs of both subjects.

Multiple statistical analyses were also performed to compare the evaluation metrics resulting from different SynX and SO cases. Paired *t*-tests were performed on RMSE and *r* values to identify significant differences in the accuracy of estimated unmeasured muscle activations between any two of the five SynX or SO cases. Paired *t*-tests were also used to identify significant differences in the accuracy of estimated muscle forces between any two of the five SynX or SO cases. In addition, paired *t*-tests were performed to compare joint moment matching errors (MAE values) between the “Params” case and the two SynX cases. All statistical analyses were performed in MATLAB with a significance level of *p* < 0.05.

## Results

### Muscle Activations

Muscle activations for unmeasured muscles estimated using SynX and SO were compared with those produced by EMG-driven model calibration using a complete set of EMG data (optimization problem “ *Params* ”). This comparison was conducted to assess the accuracy of estimated muscle activations (figure 4 and 5, table 3). Initially, during the simultaneous calibration of EMG-driven model parameters, SynX effectively estimated unmeasured muscle activations, demonstrating low RMSE values (≤ 0.17, = 0.08 ± 0.06) and moderate or strong correlation *r* values (≥0.38, = 0.55 ± 0.13) across most muscles for optimization “ *S*yn*X*_*Unmeasured*_ + *Params* ” (figure 3 and table 3). Notably, among these unmeasured muscles, SynX exhibited superior performance for the superficial muscles (e.g. rectus femoris, lateral gastrocnemius and vastus intermedius) compared to the deep-located muscles (e.g. iliacus, extensor digitorum longus and tibialis posterior) in terms of both shape and magnitude. However, the estimates for adductor muscles (RMSE = 0.01, *r ≥* 0.43) and flexor digitorum longus (RMSE = 0.05, *r* = 0.92) that typically rely on fine-wire electrodes for EMG collection remained reasonably accurate.

**Table 3.**
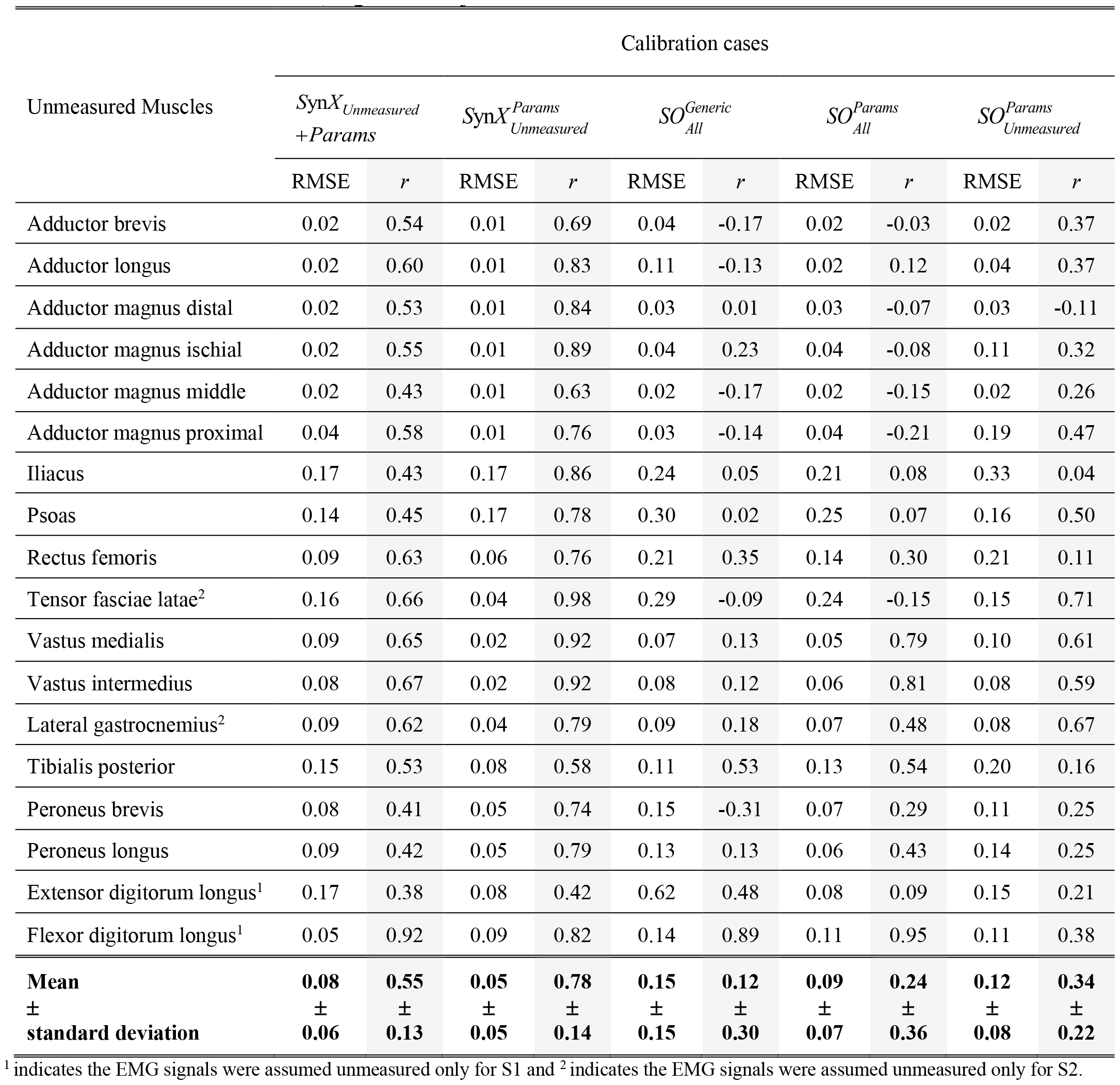
Pearson correlation coefficient r values and root mean square error (RMSE) values were calculated between the experimental (“Params” calibration) and estimated muscle activations for different calibration cases. The RMSE and *r* values for each muscle were calculated when the data across all calibration trials, legs and subjects were concatenated.

| Unmeasured Muscles | Calibration cases |  |  |  |  |  |  |  |  |  |
| --- | --- | --- | --- | --- | --- | --- | --- | --- | --- | --- |
| | $SynX_{Unmeasured}^{+Params}$ | | $SynX_{Unmeasured}^{Params}$ | | $SO_{All}^{Generic}$ | | $SO_{All}^{Params}$ | | $SO_{Unmeasured}^{Params}$ | |
| | RMSE | $r$ | RMSE | $r$ | RMSE | $r$ | RMSE | $r$ | RMSE | $r$ |
| Adductor brevis | 0.02 | 0.54 | 0.01 | 0.69 | 0.04 | -0.17 | 0.02 | -0.03 | 0.02 | 0.37 |
| Adductor longus | 0.02 | 0.60 | 0.01 | 0.83 | 0.11 | -0.13 | 0.02 | 0.12 | 0.04 | 0.37 |
| Adductor magnus distal | 0.02 | 0.53 | 0.01 | 0.84 | 0.03 | 0.01 | 0.03 | -0.07 | 0.03 | -0.11 |
| Adductor magnus ischial | 0.02 | 0.55 | 0.01 | 0.89 | 0.04 | 0.23 | 0.04 | -0.08 | 0.11 | 0.32 |
| Adductor magnus middle | 0.02 | 0.43 | 0.01 | 0.63 | 0.02 | -0.17 | 0.02 | -0.15 | 0.02 | 0.26 |
| Adductor magnus proximal | 0.04 | 0.58 | 0.01 | 0.76 | 0.03 | -0.14 | 0.04 | -0.21 | 0.19 | 0.47 |
| Iliacus | 0.17 | 0.43 | 0.17 | 0.86 | 0.24 | 0.05 | 0.21 | 0.08 | 0.33 | 0.04 |
| Psoas | 0.14 | 0.45 | 0.17 | 0.78 | 0.30 | 0.02 | 0.25 | 0.07 | 0.16 | 0.50 |
| Rectus femoris | 0.09 | 0.63 | 0.06 | 0.76 | 0.21 | 0.35 | 0.14 | 0.30 | 0.21 | 0.11 |
| Tensor fasciae latae<sup>2</sup> | 0.16 | 0.66 | 0.04 | 0.98 | 0.29 | -0.09 | 0.24 | -0.15 | 0.15 | 0.71 |
| Vastus medialis | 0.09 | 0.65 | 0.02 | 0.92 | 0.07 | 0.13 | 0.05 | 0.79 | 0.10 | 0.61 |
| Vastus intermedius | 0.08 | 0.67 | 0.02 | 0.92 | 0.08 | 0.12 | 0.06 | 0.81 | 0.08 | 0.59 |
| Lateral gastrocnemius<sup>2</sup> | 0.09 | 0.62 | 0.04 | 0.79 | 0.09 | 0.18 | 0.07 | 0.48 | 0.08 | 0.67 |
| Tibialis posterior | 0.15 | 0.53 | 0.08 | 0.58 | 0.11 | 0.53 | 0.13 | 0.54 | 0.20 | 0.16 |
| Peroneus brevis | 0.08 | 0.41 | 0.05 | 0.74 | 0.15 | -0.31 | 0.07 | 0.29 | 0.11 | 0.25 |
| Peroneus longus | 0.09 | 0.42 | 0.05 | 0.79 | 0.13 | 0.13 | 0.06 | 0.43 | 0.14 | 0.25 |
| Extensor digitorum longus<sup>1</sup> | 0.17 | 0.38 | 0.08 | 0.42 | 0.62 | 0.48 | 0.08 | 0.09 | 0.15 | 0.21 |
| Flexor digitorum longus<sup>1</sup> | 0.05 | 0.92 | 0.09 | 0.82 | 0.14 | 0.89 | 0.11 | 0.95 | 0.11 | 0.38 |
| **Mean** | **0.08** | **0.55** | **0.05** | **0.78** | **0.15** | **0.12** | **0.09** | **0.24** | **0.12** | **0.34** |
| **±** | **±** | **±** | **±** | **±** | **±** | **±** | **±** | **±** | **±** | **±** |
| **standard deviation** | **0.06** | **0.13** | **0.05** | **0.14** | **0.15** | **0.30** | **0.07** | **0.36** | **0.08** | **0.22** |

<sup>1</sup> indicates the EMG signals were assumed unmeasured only for S1 and <sup>2</sup> indicates the EMG signals were assumed unmeasured only for S2.

**Figure 4.**
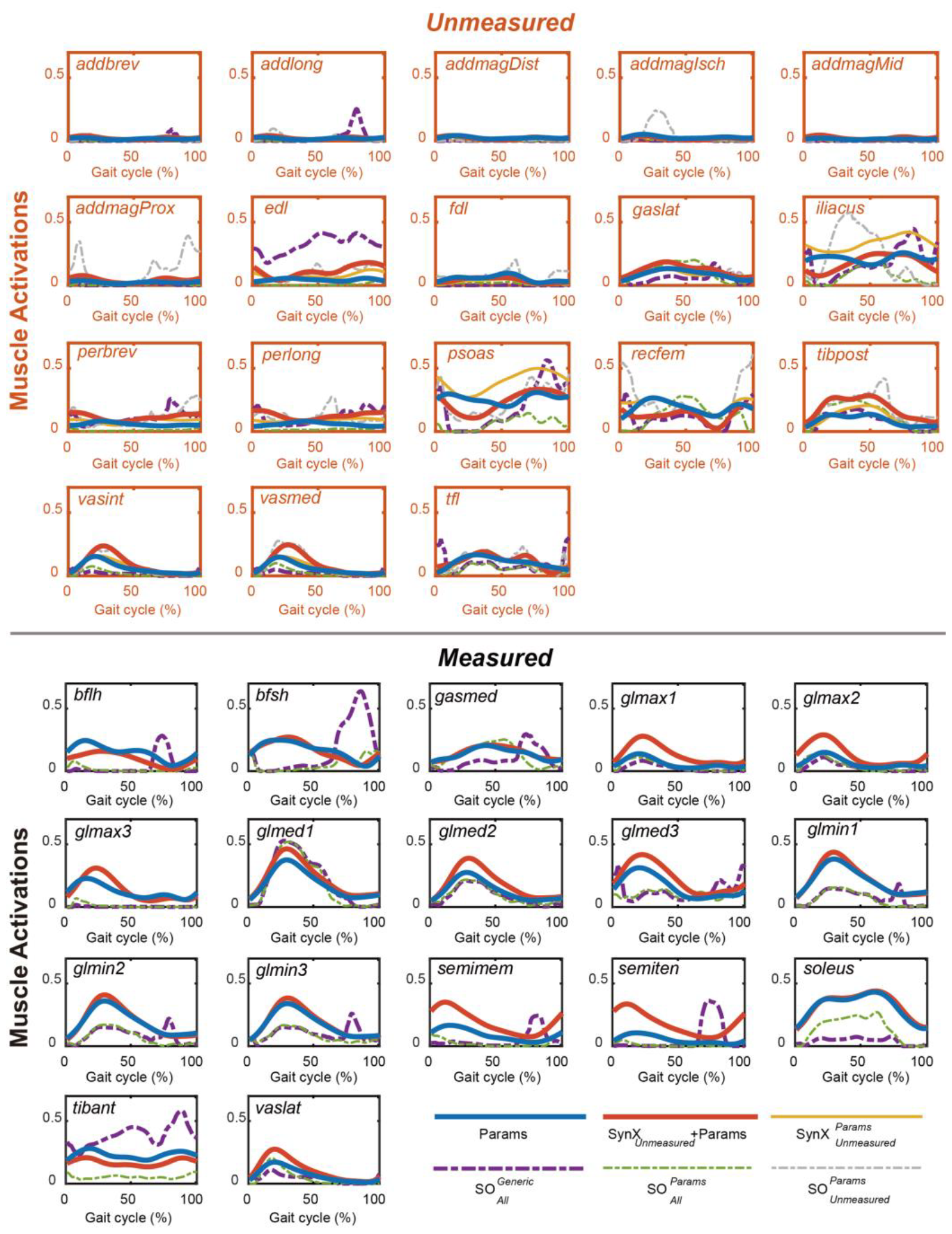
Average muscle activations for the “unmeasured” muscles (upper) and the “measured” muscles (lower) across calibration trials, legs and subjects from “*Params*” optimization (blue solid curves), SynX-based optimizations ( *S*yn*X Unmeasured* +*Params* :red solid curves and : 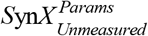 yellow solid curves and SO-based optimizations ( 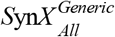: purple dash curves,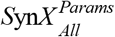: green dash curves and 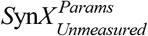 :grey dash curves). Data are reported for the complete gait cycle, where 0% indicates initial heel-strike and 100% indicates subsequent heel-strike. In addition, for the measured muscles, the curves associated with 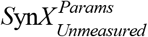 and 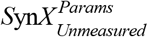 were underneath the curves associated with “*Params*” the associated muscle activations were experimental (from “*Params*” optimization) rather than calibrated.

**Figure 5.**
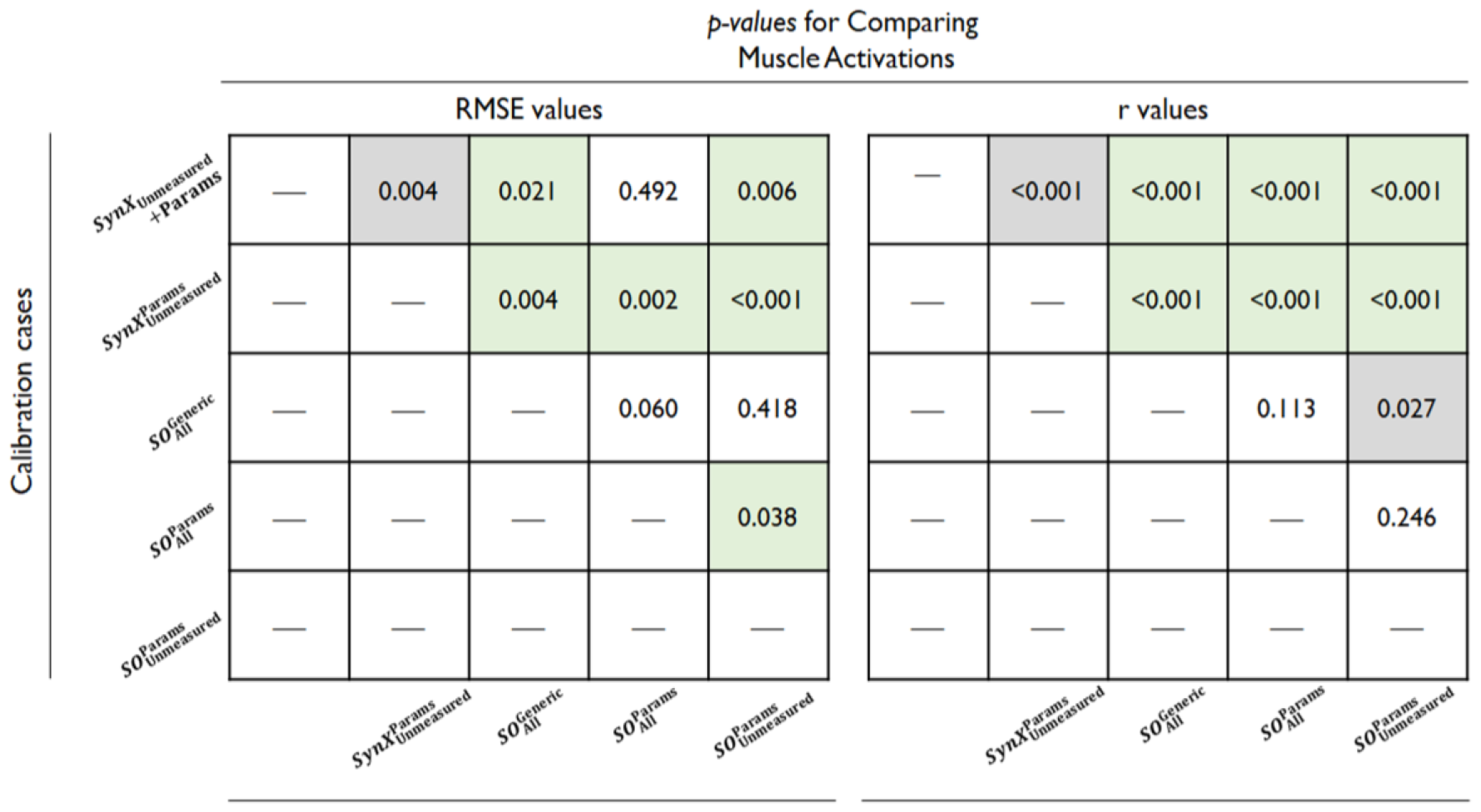
*p*-*values* obtained from paired *t-test* used to compare the estimation accuracy of muscle activations, as indicated by RMSE values (left) and *r* values (right), between different optimizations. Initially, RMSE and *r* values were calculated between the experimental (“*Params*” optimization) and estimated muscle activations from various optimizations, with the results across all calibration trials, legs, and subjects being concatenated and displayed in table 3. Subsequently, the RMSE and *r* values from each optimization were individually compared to the results from every other optimization to determine the statistical significance of the differences in estimation accuracy between each pair of optimizations. All statistical analyses were performed in MATLAB, and the level of statistical significance was set at *p* < 0.05. A box with green background indicates that the estimation performance for the y-axis optimization was significantly better (lower RMSE values or higher r values) than it for the x-axis optimization, while a box with grey background indicates that the estimation performance for the y-axis optimization was significantly worse (higher RMSE values or lower r values) than it for the x-axis optimization.

Second, SynX combined with model optimization “ *S*yn*X*_*Unmeasured*_ + *Params* ” produced significantly more accurate predictions of unmeasured muscle activations compared to the standard SO used within optimization 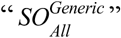. This finding was evident in terms of both magnitude (characterized by RMSE values, *p* ≤ 0.05) and shape (characterized by correlation *r* values, *p* ≤ 0.05) across unmeasured muscles and subjects (figure 4 and 5, table 3). Even for muscles with relatively low estimation accuracy using both methods, such as iliacus, psoas and extensor digitorum longus, SynX outperformed SO in reproducing the shape and magnitude of unmeasured muscle activations (table 3). Moreover, SO exhibited weak correlation (*r* ≤ 0.35) in the muscle activation predictions for the majority of unmeasured muscles within optimization 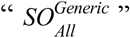, apart from tibialis posterior (*r* = 0.53), extensor digitorum longus (*r* = 0.48) and flexor digitorum longus (*r* = 0.89). Notably, the SynX-based optimization generated smooth muscle activation profiles, whereas SO exhibited discontinuities and underestimated muscle activations, featuring abrupt changes (see figure 3).

Third, both SynX-based and SO-based methods were sensitive to the level of musculoskeletal model personalization (figure 4 and 5, table 3). For SynX, employing a well-calibrated EMG-driven model for optimization 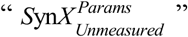 resulted in lower RMSE for the estimation of unmeasured muscle activations (RMSE = 0.05 ± 0.05) compared to optimization “ *S*yn*X*_*Unmeasured*_ + *Params* ”, when SynX variables and EMG-driven model parameter values were calibrated simultaneously. The estimated unmeasured muscle activations from optimization 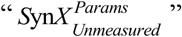 exhibited strong or very strong correlations with those generated from optimization “ *Params* ”, with the exception of the extensor digitorum longus (*r* = 0.42). For SO, well-calibrated model parameter values in optimization 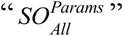 led to more accurate estimation of unmeasured muscle activations compared to using scaled generic model parameter values in optimization 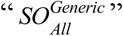, although the difference was not substantial.

Last, with the use of a well-calibrated EMG-driven model to estimate unmeasured muscle activations only, SynX in optimization case 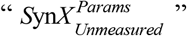 demonstrated more accurate and reliable estimates compared to SO in optimization 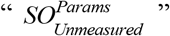. This was evidenced by remarkably lower RMSE values (*p* ≤ 0.05) and higher correlation *r* values (*p* ≤ 0.05) (figure 4 and 5, table 3). Similar to all SO-estimated muscle activations, the estimates obtained from SO in optimization 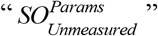 showed general underestimation and abrupt changes.

### Muscle Forces

The study quantitatively evaluated the magnitude and shape similarity of muscle forces estimated using various optimization methods and those estimated from EMG-driven modeling calibration using a full set of EMGs (optimization “ *Params* ”) through RMSE and correlation *r* measurements.

First, SynX provided reasonably accurate and reliable estimation of muscle forces that closely matched those obtained from “ *Params* ” optimization for both subjects, as shown in figure 6 and 7, table 4. In terms of unmeasured muscle forces, the RMSE values using SynX for optimization “ *S*yn*X*_*Unmeasured*_ + *Params* ” (= 101.3 ± 0.13) were significantly smaller (*p* = 0.028) than those using standard SO for optimization 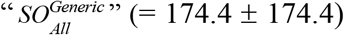. Furthermore, the correlation *r* values between the muscle forces estimated by SynX and those provided by the full EMG-driven model calibration was moderate or higher across all unmeasured muscles. Conversely, for SO, the correlation was generally weak for most muscles, except for moderate correlations observed for rectus femoris (*r* = 0.42), lateral gastrocnemius (*r* = 0.51), tibialis posterior (*r* = 0.48), and extensor digitorum longus (*r* = 0.58) (table 4).

**Table 4.**
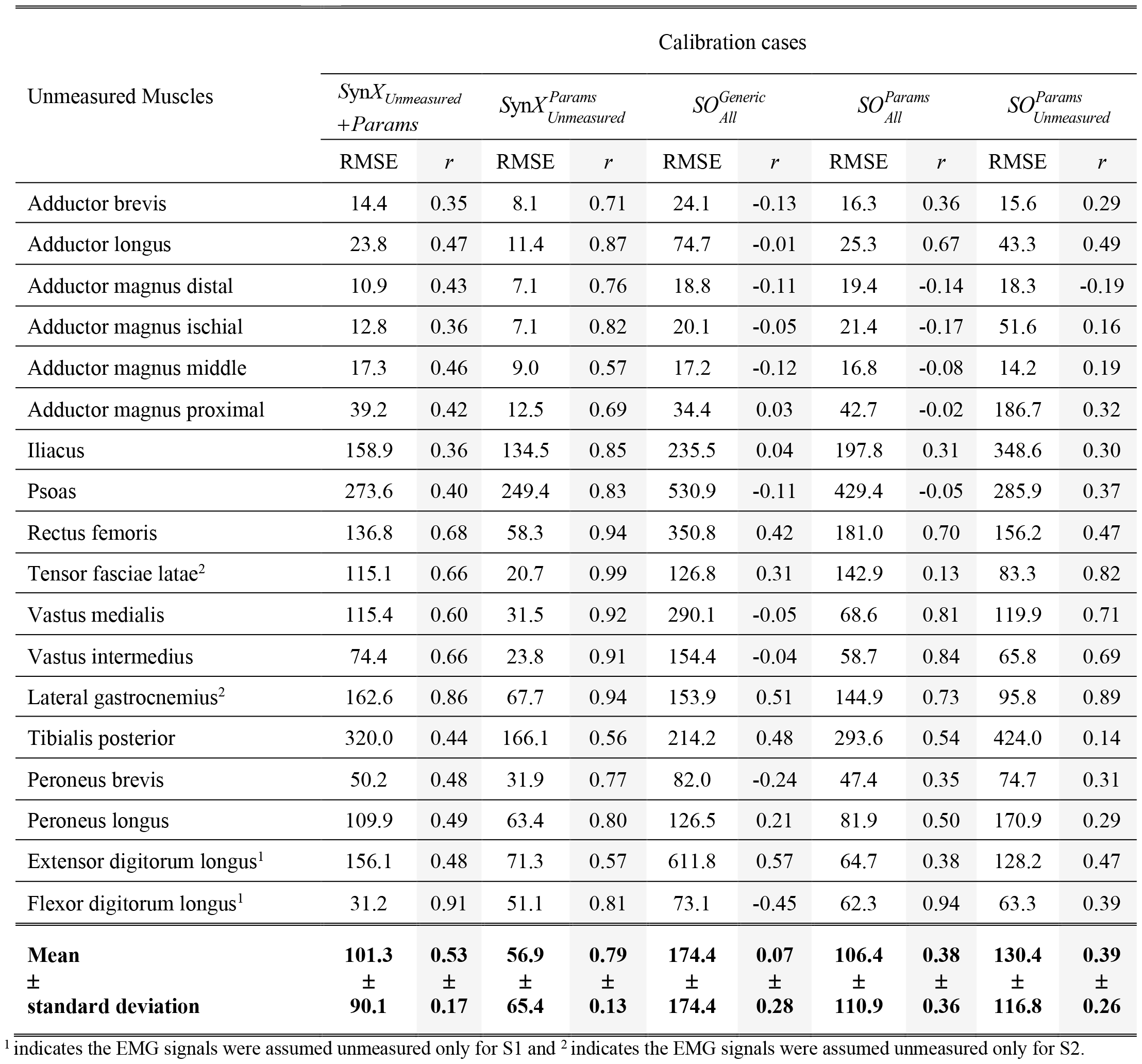
Pearson correlation coefficient r values and root mean square error (RMSE) values were calculated between the experimental (“Params” calibration) and estimated muscle forces for different calibration cases. The r and RMSE values were calculated when the data across all calibration trials and subjects were concatenated.

| Unmeasured Muscles | Calibration cases |  |  |  |  |  |  |  |  |  |
| --- | --- | --- | --- | --- | --- | --- | --- | --- | --- | --- |
| | $SynX_{Unmeasured}^{+Params}$ | | $SynX_{Unmeasured}^{Params}$ | | $SO_{All}^{Generic}$ | | $SO_{All}^{Params}$ | | $SO_{Unmeasured}^{Params}$ | |
| | RMSE | $r$ | RMSE | $r$ | RMSE | $r$ | RMSE | $r$ | RMSE | $r$ |
| Adductor brevis | 14.4 | 0.35 | 8.1 | 0.71 | 24.1 | -0.13 | 16.3 | 0.36 | 15.6 | 0.29 |
| Adductor longus | 23.8 | 0.47 | 11.4 | 0.87 | 74.7 | -0.01 | 25.3 | 0.67 | 43.3 | 0.49 |
| Adductor magnus distal | 10.9 | 0.43 | 7.1 | 0.76 | 18.8 | -0.11 | 19.4 | -0.14 | 18.3 | -0.19 |
| Adductor magnus ischial | 12.8 | 0.36 | 7.1 | 0.82 | 20.1 | -0.05 | 21.4 | -0.17 | 51.6 | 0.16 |
| Adductor magnus middle | 17.3 | 0.46 | 9.0 | 0.57 | 17.2 | -0.12 | 16.8 | -0.08 | 14.2 | 0.19 |
| Adductor magnus proximal | 39.2 | 0.42 | 12.5 | 0.69 | 34.4 | 0.03 | 42.7 | -0.02 | 186.7 | 0.32 |
| Iliacus | 158.9 | 0.36 | 134.5 | 0.85 | 235.5 | 0.04 | 197.8 | 0.31 | 348.6 | 0.30 |
| Psoas | 273.6 | 0.40 | 249.4 | 0.83 | 530.9 | -0.11 | 429.4 | -0.05 | 285.9 | 0.37 |
| Rectus femoris | 136.8 | 0.68 | 58.3 | 0.94 | 350.8 | 0.42 | 181.0 | 0.70 | 156.2 | 0.47 |
| Tensor fasciae latae<sup>2</sup> | 115.1 | 0.66 | 20.7 | 0.99 | 126.8 | 0.31 | 142.9 | 0.13 | 83.3 | 0.82 |
| Vastus medialis | 115.4 | 0.60 | 31.5 | 0.92 | 290.1 | -0.05 | 68.6 | 0.81 | 119.9 | 0.71 |
| Vastus intermedius | 74.4 | 0.66 | 23.8 | 0.91 | 154.4 | -0.04 | 58.7 | 0.84 | 65.8 | 0.69 |
| Lateral gastrocnemius<sup>2</sup> | 162.6 | 0.86 | 67.7 | 0.94 | 153.9 | 0.51 | 144.9 | 0.73 | 95.8 | 0.89 |
| Tibialis posterior | 320.0 | 0.44 | 166.1 | 0.56 | 214.2 | 0.48 | 293.6 | 0.54 | 424.0 | 0.14 |
| Peroneus brevis | 50.2 | 0.48 | 31.9 | 0.77 | 82.0 | -0.24 | 47.4 | 0.35 | 74.7 | 0.31 |
| Peroneus longus | 109.9 | 0.49 | 63.4 | 0.80 | 126.5 | 0.21 | 81.9 | 0.50 | 170.9 | 0.29 |
| Extensor digitorum longus<sup>1</sup> | 156.1 | 0.48 | 71.3 | 0.57 | 611.8 | 0.57 | 64.7 | 0.38 | 128.2 | 0.47 |
| Flexor digitorum longus<sup>1</sup> | 31.2 | 0.91 | 51.1 | 0.81 | 73.1 | -0.45 | 62.3 | 0.94 | 63.3 | 0.39 |
| **Mean** | **101.3** | **0.53** | **56.9** | **0.79** | **174.4** | **0.07** | **106.4** | **0.38** | **130.4** | **0.39** |
| **±** | **±** | **±** | **±** | **±** | **±** | **±** | **±** | **±** | **±** | **±** |
| **standard deviation** | **90.1** | **0.17** | **65.4** | **0.13** | **174.4** | **0.28** | **110.9** | **0.36** | **116.8** | **0.26** |

<sup>1</sup> indicates the EMG signals were assumed unmeasured only for S1 and <sup>2</sup> indicates the EMG signals were assumed unmeasured only for S2.

**Figure 6.**
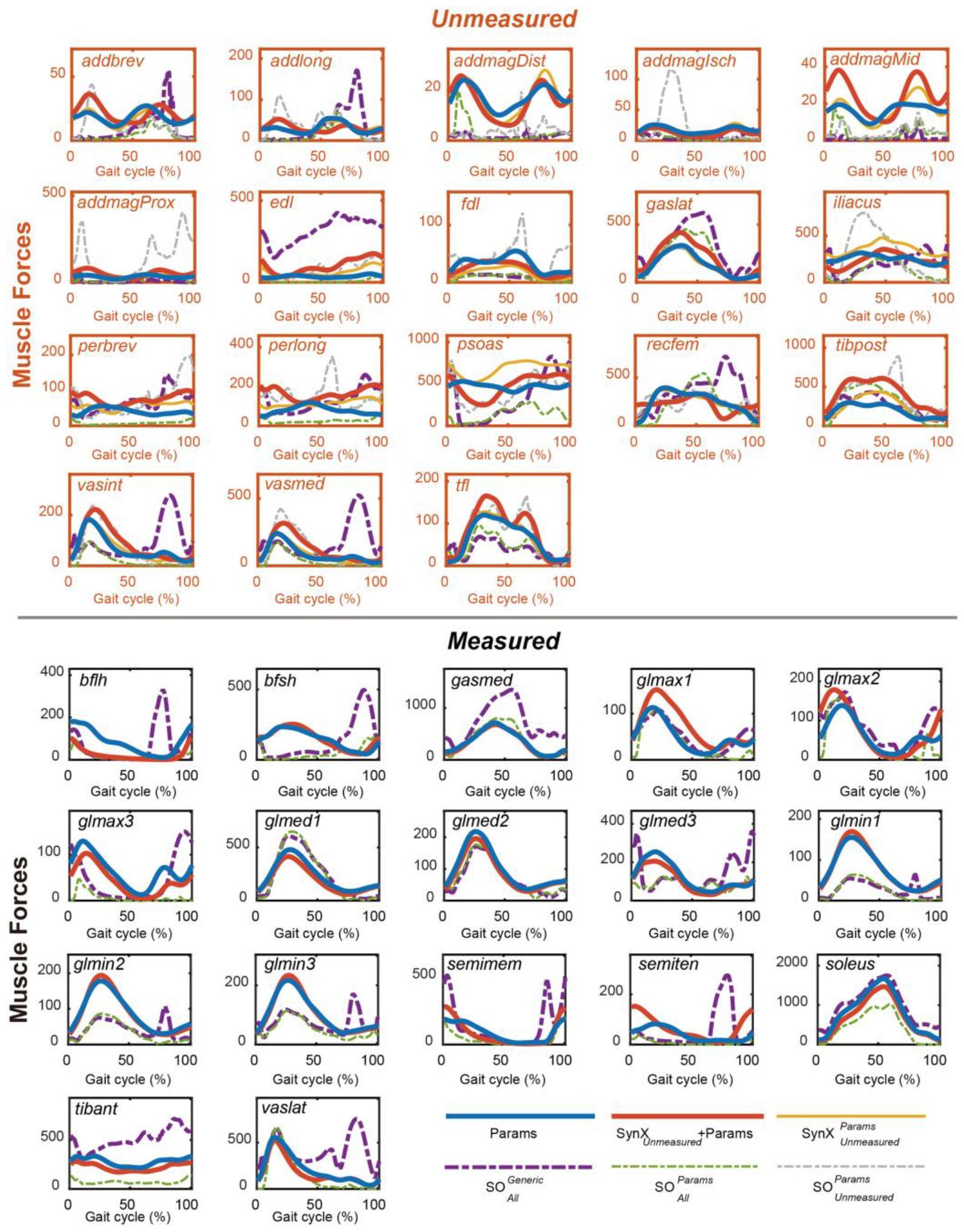
Average muscle forces for the “unmeasured” muscles (upper) and the “measured” muscles (lower) across calibration trials, legs and subjects from “*Params*” optimization (blue solid curves), SynX-based optimizations (*S*yn*XUnmeasured+* :red solid curves and 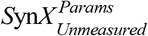: yellow solid curves and SO-based optimizations ( 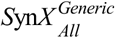 : purple dash curves, 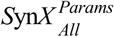: green dash curves and 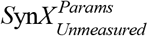 :grey dash curves). Data are reported for the complete gait cycle, where 0% indicates initial heel-strike and 100% indicates subsequent heel-strike. In addition, for the measured muscles, the curves associated with 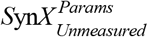 and 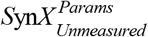 were underneath the curves associated with “*Params*” the associated muscle forces were experimental (from “*Params*” optimization) rather than calibrated.

**Figure 7.**
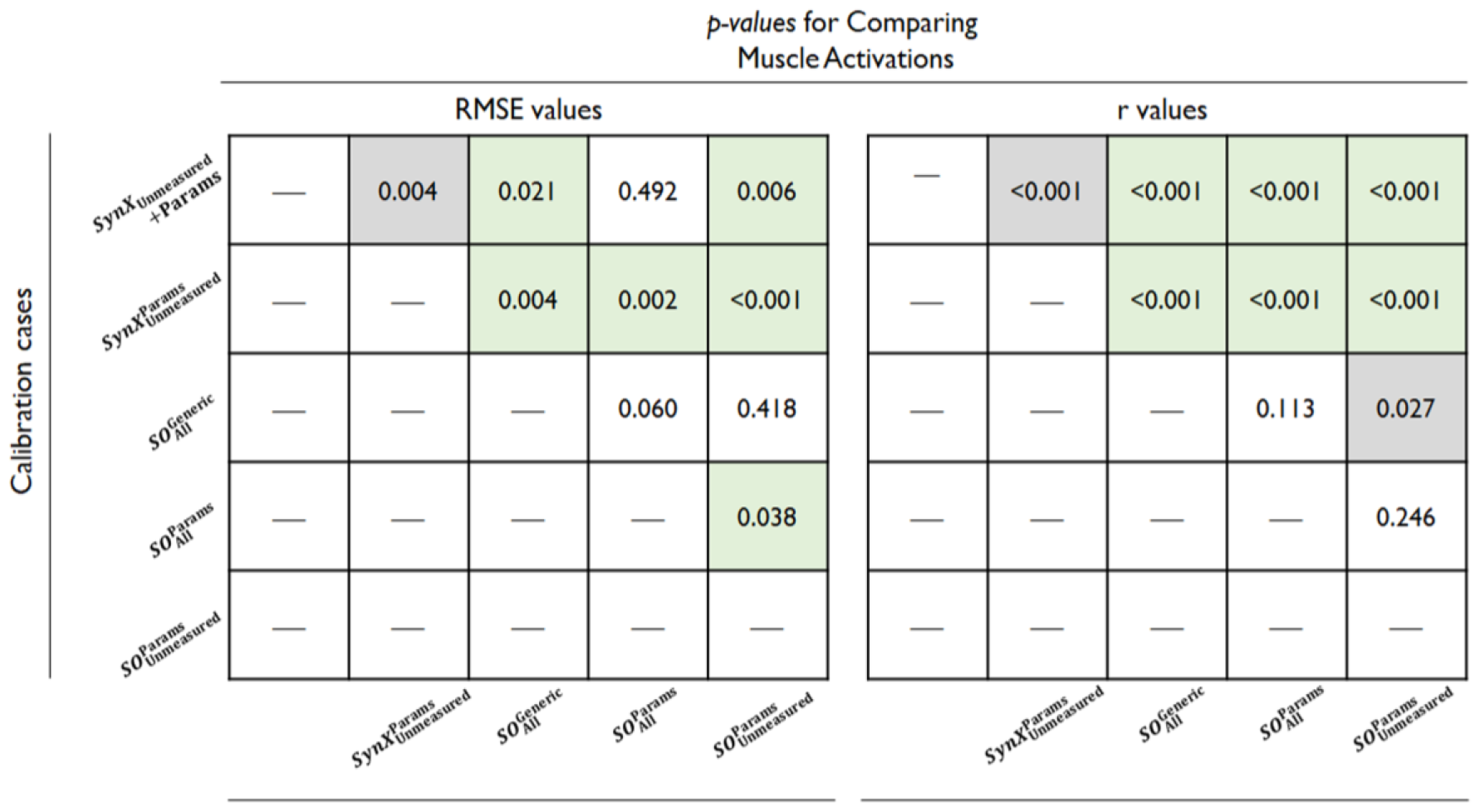
*p*-*values* obtained from paired *t-test* used to compare the estimation accuracy of muscle forces, as indicated by RMSE values (left) and *r* values (right), between different optimizations. Initially, RMSE and *r* values were calculated between the experimental (“*Params*” optimization) and estimated muscle activations from various optimizations, with the results across all calibration trials, legs, and subjects being concatenated and displayed in table 4. Subsequently, the RMSE and *r* values from each optimization were individually compared to the results from every other optimization to determine the statistical significance of the differences in estimation accuracy between each pair of optimizations. All statistical analyses were performed in MATLAB, and the level of statistical significance was set at *p* < 0.05. A box with green background indicates that the estimation performance for the y-axis optimization was significantly better (lower RMSE values or higher r values) than it for the x-axis optimization, while a box with grey background indicates that the estimation performance for the y-axis optimization was significantly worse (higher RMSE values or lower r values) than it for the x-axis optimization.

Second, model personalization had considerable influence on the accuracy of estimating muscle forces for both SynX and SO, as detailed in figure 6 and 7, and table 4. SynX demonstrated notably improved estimation accuracy in terms of both shape (*p* ≤ 0.05) and magnitude (*p* ≤ 0.05) when incorporating a well-calibrated EMG-driven model for optimization 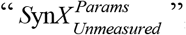, as compared to simultaneous calibration of SynX variables and EMG-driven model parameters for optimization “ *S*yn*X*_*Unmeasured*_ + *Params* ”. Similarly, SO benefited from well-calibrated model parameter values in achieving more accurate estimation of unmeasured muscle forces, leading to significantly different correlation *r* values between optimizations 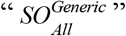 and 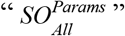, while the RMSE values remained statistically comparable between them.

Finally, with model parameter values determined through a full EMG-driven calibration, “ *Params* ”, SynX predicted unmeasured muscle forces more accurately and reliably within optimization 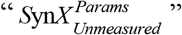 than SO within optimization 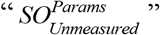. This was evidenced by significantly lower RMSE values (*p* ≤ 0.05) and higher correlation *r* values (*p* ≤ 0.05).

### Joint Moments

Compared with the MAE values between model-predicted and experimental ID joint moments from “ *Params* ” optimization, the MAE values for SynX-based optimization, with simultaneous calibration of model parameters for optimization “ *S*yn*X*_*Unmeasured*_ + *Params* ”, were consistently lower across all DoFs. On average, the MAE values were lower by 1.54 Nm for *HipFE*, 1.74 Nm for *HipAA*, 0.37 Nm for *HipRot*, 1.88 Nm for KneeFE, 2.05 Nm for *AnklePD* and 2.27 Nm *AnkleIE* across the legs of both subjects (see figure 8 and table 5). When the EMG-driven model parameter values were fixed at the values determined from “ *Params* ” optimizations, the SynX-estimated joint moments for optimization 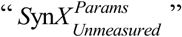 showed significantly lower MAE values only for all DoFs (*p* ≤ 0.05), except for *HipRot* (*p* = 0.078). It is important to note that inherent to the formulation of optimization, the joint moment matching errors were exceptionally small (MAEs ≤ 0.001) for all three SO-based optimizations, 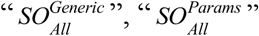, and 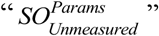.

**Table 5.** Mean absolute error (MAE) values calculated between joint moments found from inverse dynamics and either “Params” calibration or SynX-based calibrations including *S*yn*X*_*Unmeasured*_ + *Params* and 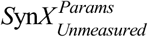

| DOFs | Calibration Case | Subjects |  |  |  |
| --- | --- | --- | --- | --- | --- |
|  |  | S1 |  | S2 |  |
|  |  | Left | Right | Left | Right |
| HipFE | *Params* | 7.45 | 6.49 | 5.81 | 4.90 |
| | $\text{SynX}_{\text{Unmeasured}} + \text{Params}$ | 4.73 | 3.96 | 3.97 | 5.84 |
| | $\text{SynX}_{\text{Unmeasured}}^{\text{Params}}$ | 8.25 | 5.55 | 6.84 | 7.53 |
| HipAA | *Params* | 7.31 | 7.23 | 7.17 | 5.51 |
| | $\text{SynX}_{\text{Unmeasured}} + \text{Params}$ | 4.88 | 5.65 | 4.85 | 4.90 |
| | $\text{SynX}_{\text{Unmeasured}}^{\text{Params}}$ | 5.79 | 5.80 | 6.42 | 2.77 |
| HipRot | *Params* | 5.82 | 5.64 | 2.53 | 2.19 |
| | $\text{SynX}_{\text{Unmeasured}} + \text{Params}$ | 5.61 | 4.81 | 2.55 | 1.75 |
| | $\text{SynX}_{\text{Unmeasured}}^{\text{Params}}$ | 5.98 | 5.22 | 2.87 | 1.46 |
| KneeFE | *Params* | 6.33 | 4.26 | 4.90 | 4.70 |
| | $\text{SynX}_{\text{Unmeasured}} + \text{Params}$ | 3.98 | 2.68 | 3.08 | 2.92 |
| | $\text{SynX}_{\text{Unmeasured}}^{\text{Params}}$ | 3.94 | 2.87 | 3.88 | 2.02 |
| AnklePD | *Params* | 6.88 | 5.68 | 5.04 | 4.85 |
| | $\text{SynX}_{\text{Unmeasured}} + \text{Params}$ | 4.51 | 4.11 | 3.14 | 2.50 |
| | $\text{SynX}_{\text{Unmeasured}}^{\text{Params}}$ | 5.78 | 4.63 | 4.14 | 1.77 |
| AnkleIE | *Params* | 4.89 | 2.82 | 8.19 | 1.75 |
| | $\text{SynX}_{\text{Unmeasured}} + \text{Params}$ | 2.50 | 1.89 | 2.73 | 1.46 |
| | $\text{SynX}_{\text{Unmeasured}}^{\text{Params}}$ | 2.90 | 2.13 | 4.22 | 0.95 |

**Figure 8.**
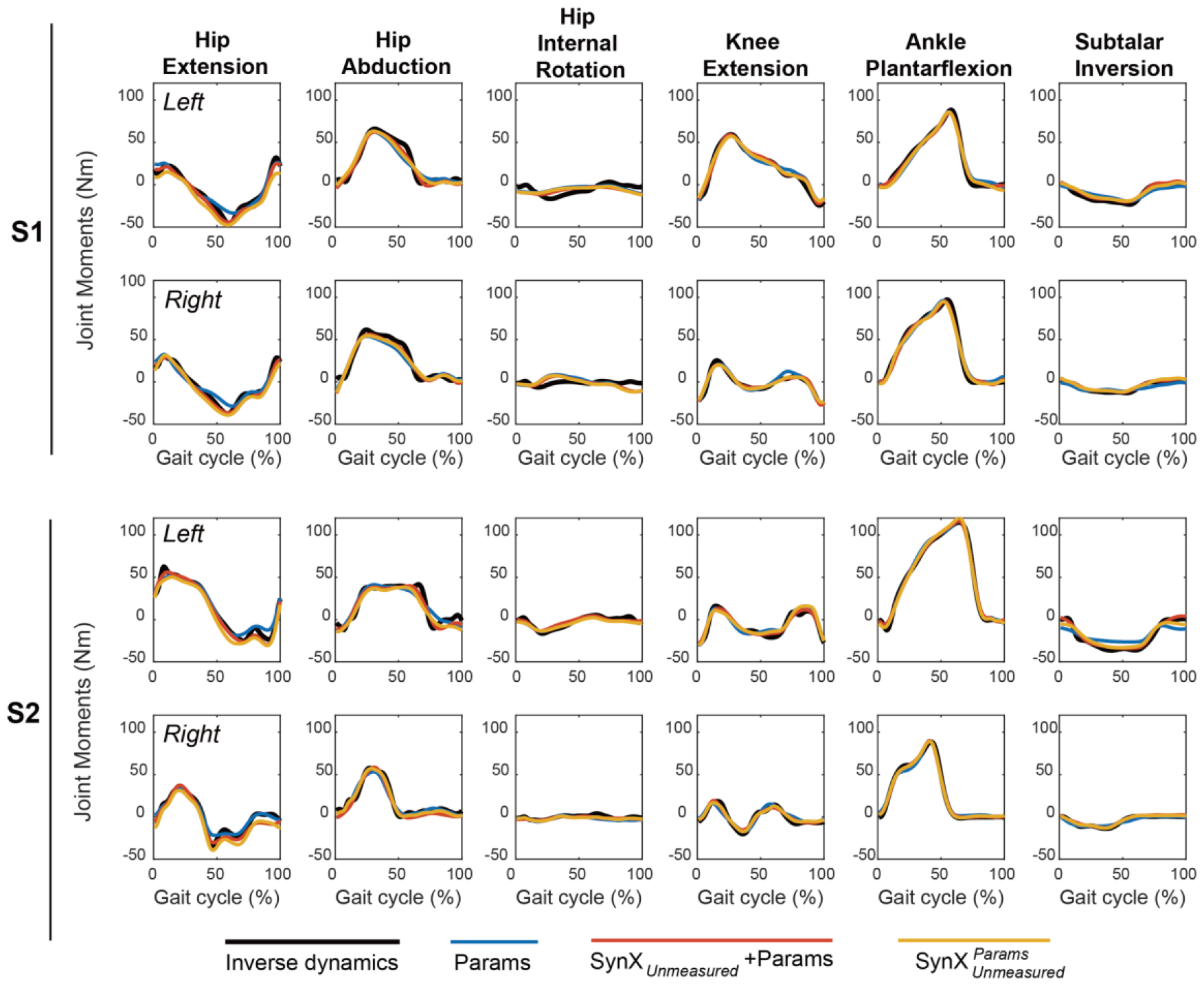
Average joint moments across calibration trials from “*Params*” optimization (blue solid curves) and SynX-based optimizations ( *S*yn*X*_*Unmeasured*_ + *Params*: red solid curves and 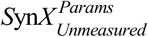: yellow solid curves). Data are reported for the complete gait cycle, where 0% indicates initial heel-strike and 100% indicates subsequent heel-strike.

### Model Parameters

In general, the four activation dynamic model parameters and the two Hill-type muscle-tendon model parameters from optimization showed a high degree of similarity between “*S*yn*X*_*Unmeasured*_ + *Params* ” optimization and “ *Params* ” optimization for the measured muscles (refer to figure 9, left panel). Conversely, for the unmeasured muscles, when simultaneously tuning SynX variables, “*S*yn*X*_*Unmeasured*_ + *Params* ” still maintained the pattern defined by the parameter magnitudes of the optimal fiber length and tendon slack length for each model parameter (refer to figure 9, right panel). However, substantial discrepancies in the four activation dynamic model parameters were observed for the unmeasured muscles between the SynX approach for “*S*yn*X*_*Unmeasured*_ + *Params* ” and the full EMG-driven model calibration for “ *Params* ”.

**Figure 9.**
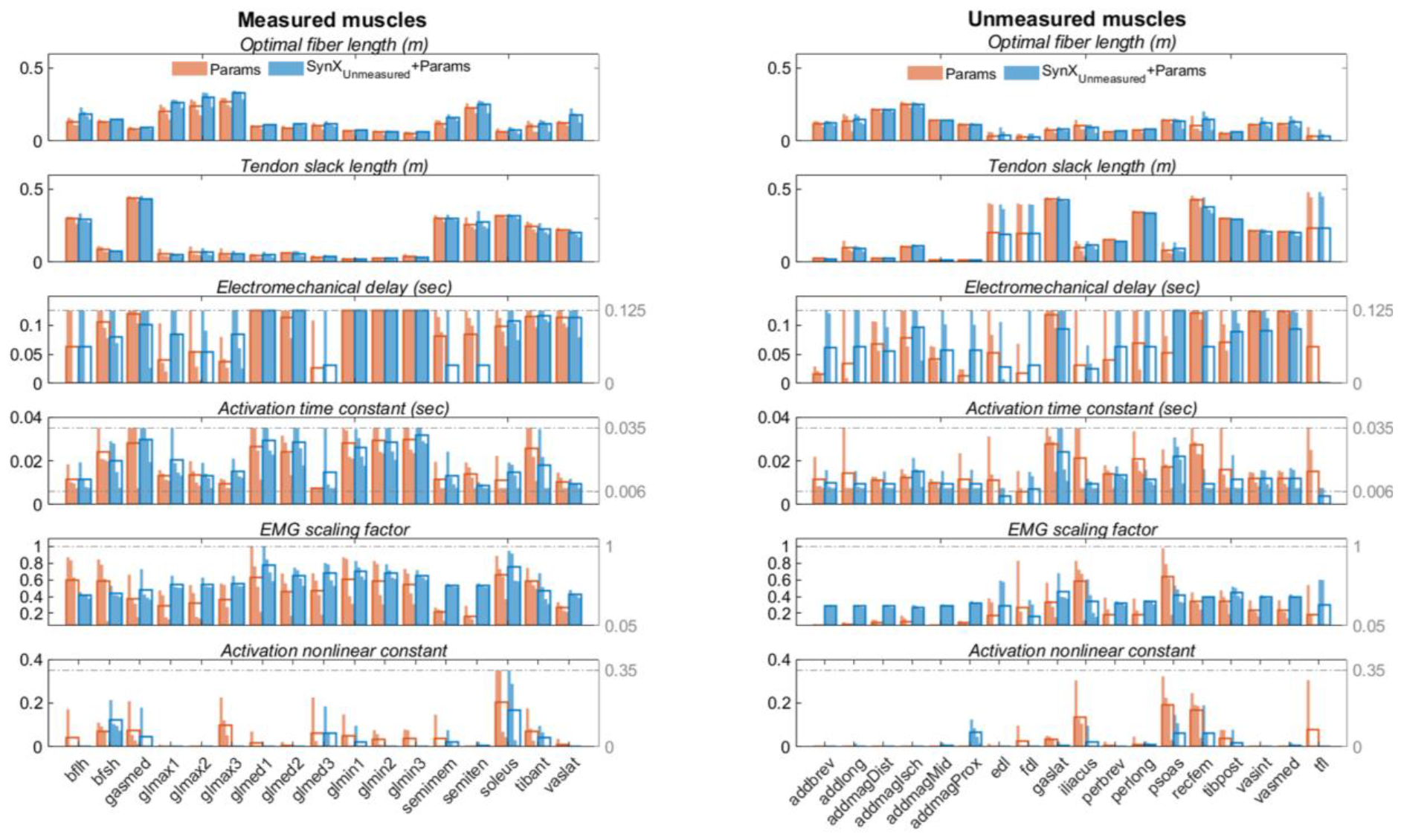
EMG-driven model parameters of two legs of both subjects from “gold standard (*Params*)” optimization (in blue) and “*S*yn*X*_*Unmeasured*_ + *Params* ” optimization (in orange). The upper and lower bounds for each of the four activation dynamics model parameters during optimization have been indicated by grey dash-sot lines, where the upper and lower bounds for the scaling factors of optimal fiber lengths and tendon slack lengths were [0.6, 1.4] for all muscles.

## Discussion

This study extended the capability of synergy extrapolation (SynX) to enable the concurrent estimation of a large number of unmeasured muscle excitations and calibration of an EMG-driven model. The approach was developed and evaluated using gait datasets collected from two post-stroke subjects performing treadmill walking at self-selected and fastest-comfortable speeds. EMG signals measured bilaterally from eight muscles were treated as “unmeasured” and estimated using the synergy information extracted from another eight muscles treated as “measured.” The muscle activations, forces, and model parameter values for the unmeasured muscles were quantitatively compared to “gold standard” values obtained when all 16 channels of EMG data were used to calibrate an EMG-driven musculoskeletal model for each leg of each subject. The results revealed that the estimated unmeasured muscle activations and forces were reasonably accurate and reliable in term of both shaped and magnitude (figures 4 and 6, Tables 3 and 4). Moreover, Hill-type muscle-tendon model parameter values for both unmeasured and measured muscles, including optimal fiber length and tendon slack length, exhibited a high level of agreement with the “gold standard” model parameter values (figure 9). When SO estimates of unmeasured muscle activations and forces were compared with SynX estimates, the SynX results were more accurate and realistic than those from SO (figures 4 and 6, tables 3 and 4), which contained abrupt changes and tended to underestimate the unmeasured muscle quantities. When the sensitivity of estimated unmeasured muscle activations and forces to the level of model personalization was investigated, both SynX and SO generated substantially more accurate estimates when utilizing well-calibrated muscle-tendon parameters. However, SynX demonstrated superior performance to SO in estimating unmeasured muscle activations and forces when employing model parameter values from full EMG-driven model calibration.

SynX has demonstrated superior performance over SO for estimating muscle activations and forces for several important reasons. First, by utilizing measured synergy excitations as building blocks, it reduced the problem of finding unknown time-varying muscle excitations to identifying a small number of unmeasured synergy vector weights. This led to a substantial reduction in the search space for the optimization in comparison with SO-based approaches[37]. Second, unlike SO-based approach, which solved a time frame of muscle activation at a time, the inherent constraints of dependence between time frames in weighted synergy excitations resulted in smooth and continuous estimated muscle activations, improving the physiological plausibility of the estimates. Third, the time-invariance of unmeasured and residual synergy vector weights enabled a single-layer optimization process, simultaneously achieving EMG-driven model personalization and muscle activation estimation, which enhanced the accuracy of muscle force estimation. Fourth, calibration of synergy-structured residual muscle excitations was integrated into SynX to enhance the accuracy of predicted unmeasured muscle excitations. Unlike SO, where unmeasured muscle activations, as design variables, inclined to deviate from experimental muscle excitations during iterative adjustments for minimizing joint moment matching errors, SynX introduced residual muscle excitations to account for joint moment matching errors, preventing predicted missing muscle excitations from excessively compensating for joint moment prediction inaccuracy through optimization. The addition of residual muscle excitations in turn prevented inaccuracy as a consequence[38]. Fifth, the SynX-based methods did not require assumptions in the optimization process, whereas SO approach led to an underestimation of muscle activations by minimizing co-activation between agonist and antagonist muscles concurrently[30,53]. Last, it has been theorized that muscle synergies are generated by the central nervous system to efficiently regulate the control of highly redundant musculoskeletal systems [54,55]. The SynX-based approach leveraged the concept of muscle synergy, making the method more physiological reasonable.

SynX offers benefits over other computational methods for the estimating missing EMG signals within [25,56–58] or outside [59–61] the context of musculoskeletal modeling. Below are some representative approaches that offer great insights for us to develop our method. First, one such method utilizes Gaussian process regression models to describe the synergistic relationship between a subset of muscles, which enables the estimation of unmeasured muscle excitations using information provided by a subset of measured muscle excitations [60]. However, the muscle excitations associated with “unmeasured” muscles must be initially known for conducting the required model training process, rendering this method infeasible when the “unmeasured” muscle excitations are truly unmeasurable due to experimental constraints or safety considerations. Second, an alternative approach employs low-dimension sets of impulsive excitation primitives to estimate unmeasured muscle excitations [25,56,57]. Each muscle is assigned to a module by evaluating associated weighting factors for the excitation primitives derived from measured muscle excitations. Muscles without EMG signals are assumed to belong to the same module as measured muscles that share the same innervation and contribute to the same mechanical action. Meanwhile, the primitive-driven excitations for measured muscles are minimally adjusted to improve joint moment estimation in EMG-assisted models. However, these adjustments masked the omission of active force generating properties for unmeasured muscles (i.e., iliacus and psoas), resulting in noticeable hip joint moment prediction errors. Furthermore, none of these studies evaluated the accuracy of predicted unmeasured muscle excitations due to the lack of corresponding experimental EMG data. Third, hybrid EMG-informed models that incorporates SO to determine unmeasured muscle activations have been developed [28,29]. Satori et al. also allowed minimal adjustments of measured muscle activations while predicting missing EMG signals (e.g., from iliacus and psoas) using SO[28]. However, none of these methods have provided evidence that estimation of unmeasured muscle activations was reliable and in reasonable agreement with experimental measurements. Furthermore, due to the nature of SO, the resulting muscle activations might exhibit unrealistic discontinuities. Last, as another well-established approach within OpenSim, the computed muscle control (CMC) algorithm solves a static optimization to determine muscle excitations necessary for achieving the desired accelerations for tracking experimental motion, providing more accurate joint moments compared to SO [58,62– 64]. However, it has been observed that CMC may be less robust and computationally efficient when estimating muscle function in human locomotion. All in all, EMG-driven modeling method with SynX provides an enhanced approach for estimating unmeasured muscle excitations, forces and joint moments in an efficient manner, without the requirement for prior knowledge of the “unmeasured” muscle excitations during the model training phase.

This study quantified the impact of model personalization, specifically focusing on muscle-tendon parameters, on the estimation of muscle activations and forces in both EMG-driven modeling with SynX and SO. In the case of SynX, the tracking errors between the estimated and experimental estimates were remarkably reduced when muscle-tendon parameter values were personalized to a suitable level for optimization 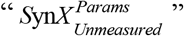. Meanwhile, the mean correlations between the estimated and experimental values were also substantially increased, moving from moderate to strong. Additionally, the matching errors of joint moments during optimization 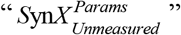 stayed closer to those obtained from full EMG-driven model optimization “ *Params* ”, as opposed to calibrating model parameters concurrently within optimization “ *S*yn*X*_*Unmeasured*_ + *Params* ”. In the case of SO, consistent with previous studies [65], personalization of muscle-tendon parameters showed noticeable improvements in estimating of muscle activations and forces in terms of both shape and amplitude for optimization 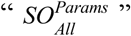, with statistically significant enhancement observed only for the shape of muscle forces. In light of these observations, enhancing the level of model personalization generally improved the accuracy of estimation. However, there were significant variations in the degree of improvement among different approaches. In scenarios where a well-calibrated musculoskeletal model is available, SynX has the ability to predict muscle activations for muscles lacking EMG data with reasonable amplitude and shape, whereas SO can predict unmeasured muscle activations with reasonable amplitude but not accurate shape. When conducting simulations using a scaled generic model, SynX successfully replicated muscle activations with the correct amplitude and shape, which SO did not achieve this.

Joint moment matching errors differ among optimizations using different approaches (figure 8 and table 5). First, the inverse dynamics (ID) and estimated joint moments exhibited a much closer agreement in the SO-based optimizations, 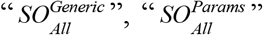, and 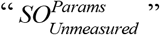 than in the EMG-driven modeling optimizations, “ *Params* ”, “ *S*yn*X*_*Unmeasured*_ + *Params* ”, and 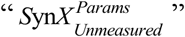. The optimization formulation used by SO in equation (10) resulted in extremely small joint moment matching errors. However, the additional constraints within EMG-modeling methods, including muscle activation-contraction dynamics and the dependency between time frames of EMG signals, limited the torque-generating capacity of muscles, thereby preventing the reproduction of joint moments. Second, the joint moment matching errors, arranged in descending order, for optimizations associated with the EMG-driven modeling method are “ *Params* ”,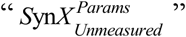 and “ *S*yn*X*_*Unmeasured*_ + *Params* ”. This observation was attributed to the increasing degrees of freedom in the optimization, determined by additional SynX variables for 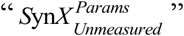, and also additional model parameters for “ *S*yn*X*_*Unmeasured*_ + *Params* ”, which enabled the optimizer to reduce the joint moment matching errors. This can also explain why the joint moment matching errors were smaller when estimating 8 channels of unmeasured EMG signals in this study compared to when estimating EMGs for only the iliacus and psoas in [38]. Last, regardless of the level of model personalization and the number of channels of muscle activations to estimate, SO consistently found the solutions of muscle activations at each time frame to almost perfectly match the ID joint moments, although occasionally requiring a small amount of reserve actuator torque due to model inadequacies. Consequently, static optimization does not possess the joint moment matching errors needed to calibrate muscle-tendon model parameters.

This study also examined the impact of incorporating the SynX process in the EMG-driven modeling framework on the calibrated model parameter values. It was observed that Hill-type muscle-tendon model parameter values, specifically optimal fiber length and tendon slack length, from optimization “ *S*yn*X*_*Unmeasured*_ + *Params*” closely approximated the “true” values obtained from full EMG-driven model optimization “ *Params* ”, as depicted in figure 9. However, the activation dynamics model parameters, including electromechanical delay, activation time constant, EMG scale factor, and activation nonlinear constant, were only reproduced with reasonable similarity for the measured muscles. When SynX was used to estimate a large number of missing EMGs during EMG-driven model calibration, it introduced additional flexibility through SynX variables into the optimization problem. This, in turn, had a cascading effect on the calibrated model parameter values for all muscles. Thus, beyond the primary cost terms specified in equation (9), penalty terms, functioning as ”soft constraints” were incorporated to restrict deviations of model parameter values from the initial model or a designated reference value[23,37,38]. The objective was to minimize the impact of SynX on model parameter values. The strategy proved effective for the Hill-type muscle-tendon models, which are inherently highly nonlinear. Nevertheless, it remained challenging to maintain the values of the activation dynamics model parameters through the utilization of penalty terms. Typically, the transformations from muscle excitations to muscle activations were determined by activation dynamics model parameters, such as time shifts typically defined by electromechanical delay and amplitude scaling dictated by EMG scale factors. Within SynX, however, unmeasured muscle excitations were constructed using linear combinations of measured synergy excitations, which could already account for these potential transformations. Consequently, numerous combinations of model parameters values and SynX variables could result in identical muscle activations. Thus, without compromising the accuracy of muscle activations, the activation model parameter values might approach the designated values in the penalty terms, allowing SynX variables to adjust to provide required muscle activations.

The inclusion of the SynX in the EMG-driven model calibration process had minimal impact on the estimation of measured muscle activations and forces, as depicted in figures 4 and 6, respectively. The muscle activations and forces estimated by SynX for measured muscles from optimization “ *S*yn*X*_*Unmeasured*_ + *Params* ” remained closely aligned with those from calibration “ *Params* ”, in contrast to the results from the commonly formulated 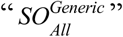 optimization. While the most significant deviations in the SynX-estimated muscle activations occurred for semimembranosus and semitendinosus (figure 4), and the greatest discrepancies in the SynX-estimated muscle forces were observed for the muscles spanning the hip joint, such as biceps femoris long head and gluteus maximus superior (figure 6), they still exhibit a similar shape to the experimental curves. These observations were largely attributed to the optimization formulation for the SynX-incorporated EMG-driven calibration, resulting in minimal changes in the calibrated activation dynamics model and Hill-type muscle-tendon model parameter values (figure 9).

Several methodological choices for SynX were necessary to consider, as they could potentially impact SynX performance, as indicated in table 2. A series of previously published studies from the author have extensively investigated various methodological choices, with the goal of identifying an optimal combination that could yield the most reliable and accurate estimation of unmeasured muscle activations [36–38]. Initially, principal component analysis (PCA) provided more accurate, reliable, and efficient estimates of unmeasured muscle excitations compared to non-negative matrix factorization (NMF), due to the non-negativity constraints for NMF and extra design variables for PCA, both of which could result in a more restricted feasible search space for NMF in comparison to PCA [37,38]. Additionally, PCA was particularly beneficial in our framework because it permitted residual excitations to be both positive and negative, which could be beneficial for achieving lower joint moment errors. Second, by comparing the results of five different EMG normalization methods that were performed either within individual trials or across all trials, we observed that EMG normalization does not have a significant influence on the SynX performance[37]. As a result, the measured muscle excitations were normalized to their maximum values across all trials before MSA to facilitate easy implementation. Furthermore, as the number of synergies increased, the performance of SynX exhibited non-monotonic behavior, with five and six synergies generally providing the best SynX performance and outcomes for EMG-driven model calibration[38]. Hence, when an increasing number of muscles were treated “unmeasured” in this study, five synergies were selected for generating the results in this study, considering the reduction of computational costs. Last, according to the assumptions about the variability of synergy vector weight across walking trials, we categorized them associated with unmeasured and residual muscle excitations as trial-specific, speed-specific, and subject-specific, respectively, while different concatenation strategies were used to extract corresponding synergy excitations. it was indicated that with an equal number of synergies, the trial-specific unmeasured synergy vector weights and speed-specific residual synergy vector weights produced the best SynX performance for the majority of subjects[38]. This insight shed lights on the categorization strategy of synergy vector weight across walking trials within this study.

A reasonable choice of a neural control strategy is essential for producing physiologically realistic predictive simulations of walking [66]. To date, prevailing predictive simulation studies have explored the optimality of neural control principles underlying human gait, and commonly reported that minimizing the sum of squares of muscle activations in the cost function, a typical practice in SO, can result in a human-like walking pattern [26,66,67]. However, the comparative results between the estimated muscle activations and forces from both SynX and SO in this study have raised a pivotal question: If the prevalent neural control strategy of minimizing the sum of squares of muscle activations fails to accurately estimate muscle activations when the joint kinematics and moments are known a priori from experimental walking data, how can it provide reliable estimates in the predictive simulations of walking when the joint kinematics and moments are unknown a priori? These findings may also yield valuable insights into the potential benefits of muscle synergies for predicting walking motion with musculoskeletal models. While the reliability of a synergy-based neural control strategy for generating predictive simulations of walking has been preliminary verified for only one experimental scenario thus far [68], the results of the present study endorse further exploration of a synergy-based neural control strategy for generating predictive simulations of walking.

This study exhibited several limitations which may provide insights for future research endeavors. First, this study validated the effectiveness of our EMG-driven modeling framework incorporating SynX by analyzing gait datasets from two post-stroke subjects, as these experimental datasets provided EMG signals for every muscle in our musculoskeletal model, enabling the evaluation of estimation accuracy. Further investigation is necessary to investigate diverse subject populations with larger sample sizes. Second, we developed the framework using walking data with two representative speeds. It would be valuable to investigate its applicability for various dynamic movement conditions and experimental scenarios, including stair climbing and running. Third, to enhance computational efficiency, we integrated a rigid tendon model into our Hill-type muscle-tendon models. Research studies have indicated that rigid and compliant tendon models produce almost identical muscle force estimates for slow movements like walking at a healthy speed, but different muscle force estimates for faster movements such as running [69,70]. As both of our stroke subjects walked at slow speeds, it suggests that use of a rigid tendon model was appropriate. However, it would be worthwhile to expand our approach to include compliant tendons in our Hill-type muscle-tendon models, enabling the applications to tasks involving fast movements. Last, we analyzed the impact of personalizing muscle-tendon parameter values on SynX performance. However, various other aspects of model personalization, including skeletal geometries, muscle kinematics, and other physiological properties that contribute to muscle force generation, may also impact muscle force estimates. Future work should therefore aim to extend the methods to investigate whether SynX performance was sensitive to these aspects of model personalization. The author has recently developed an EMG-driven modeling method that can personalize muscle wrapping surface parameters [71]. Therefore, one of the forthcoming research directions would focus on examining the influence of personalizing muscle-tendon pathway on SynX-based estimates.

## Conclusions

In conclusion, this study demonstrated a significant advancement over previous research by highlighting the capability of SynX to reproduce a large number of unmeasured muscle excitations while simultaneously calibrating EMG-driven model parameter values. Notably, the estimation accuracy of muscle activations and forces in terms of shape and amplitude for the unmeasured muscles, was significantly higher than that of the standard SO approach. The incorporation of SynX process had minimal impact on the calibrated Hill-type muscle-tendon model parameter values for all muscles and activation dynamics model parameter values for the measured muscles. Additionally, when integrated with well-calibrated musculoskeletal models, both SynX and SO produced substantially more accurate estimates of unmeasured muscle activations and forces, with SynX demonstrating superior performance over SO in this regard. The findings suggest that SynX could effectively address the practical challenge of collecting a full set of EMG signals for EMG-driven modeling calibration in the lower extremity during walking, with significant implications for personalized treatments for muscle impairments in situations where difficulties arise in collecting EMG signals from all important contributing muscles.

## Declarations

### Ethics approval and consent to participate

All experimental procedures were performed in accordance with Declaration of Helsinki and approved by the University of Florida Health Science Center Institutional Review Board (IRB-01), and the subject provided written informed consent before participation.

### Consent for publication

Not applicable.

### Availability of data and materials

The SynX EMG-driven modeling process presented in this study is freely available within the Muscle-tendon Model Personalization Tool provided with the open-source Neuromusculoskeletal Modeling Pipeline software (https://nmsm.rice.edu).

### Authors’ contributions

B.J.F. designed and performed the experiments; D.A. wrote the programs; D.A. analyzed the data, prepared figures, and drafted the manuscript; D.A. wrote the manuscript; D.A. and B.J.F. revised the manuscript; D.A. and B.J.F. approved the final version of the manuscript.

## Acknowledgements

Not applicable.

## List of abbreviations

EMG: electromyography
SynX: synergy extrapolation
SO: static optimization
RMSE: root mean square errors
DOFs: degrees of freedom
IK: inverse kinematics
ID: inverse dynamic
MSA: muscle synergy analysis
PCA: principal component analysis
NMF: non-negative matrix factorization

